# Fibroblasts-derived from Pluripotent Cells Harboring a Single Allele Knockout in Two Pluripotency Genes Exhibit DNA Methylation Abnormalities and pluripotency induction Defects

**DOI:** 10.1101/2022.05.18.492474

**Authors:** Rachel Lasry, Noam Maoz, Albert W. Cheng, Nataly Yom Tov, Elisabeth Kulenkampff, Meir Azagury, Hui Yang, Cora Ople, Styliani Markoulaki, Dina A. Faddah, Kirill Makedonski, Ofra Sabbag, Rudolf Jaenisch, Yosef Buganim

## Abstract

A complete knockout (KO) of a single key pluripotency gene has been shown to drastically affect embryonic stem cell (ESC) function and epigenetic reprogramming. However, knockin (KI)/KO of a reporter gene only in one of two alleles in a single pluripotency gene is considered harmless and is largely used in the stem cell field. Here, we sought to understand the impact of simultaneous elimination of a single allele in two ESC key genes on pluripotency potential and acquisition. We established multiple pluripotency systems harboring KI/KO in a single allele of two different pluripotency genes (i.e. Nanog^+/-^; Sall4^+/-^, Nanog^+/-^; Utf1^+/-^, Nanog^+/-^; Esrrb^+/-^ and Sox2^+/-^; Sall4^+/-^). Interestingly, although these double heterozygous mutant lines maintain their stemness and contribute to chimeras equally to their parental control cells, fibroblasts derived from these systems show a significant reduction in their capability to induce pluripotency either by Oct4, Sox2, Klf4 and Myc (OSKM) or by nuclear transfer (NT). Tracing the expression of Sall4 and Nanog, as representative key pluripotency targeted genes, at early phases of reprogramming could not explain the seen delay/blockage. Further exploration identifies abnormal methylation landscape around pluripotent and developmental genes in the double heterozygous mutant fibroblasts. Accordingly, treatment with 5-azacytidine two days prior to transgene induction rescues the reprogramming defects. This study emphasizes the importance of maintaining two intact alleles for pluripotency induction and suggests that insufficient levels of key pluripotency genes leads to DNA methylation abnormalities in the derived-somatic cells later on in development.

## INTRODUCTION

Fluorescent reporter genes are widely used to monitor cell states such as stemness, differentiation, cell cycle and migration (Benchetrit et al., 2019; Eastman et al., 2020). One common strategy to introduce a reporter gene is by a knockin/knockout (KI/KO) approach. In this strategy, a fluorescent gene is introduced into a *locus* of interest by replacing the endogenous gene with a fluorescent reporter, leaving the targeted gene with only one functional allele. Many fluorescent reporter cell lines have been generated over the years using this approach, targeting either pluripotency genes such as *Sox2* (Arnold et al., 2011; Avilion et al., 2003), *Nanog* (Meissner et al., 2007; Wernig et al., 2008) and *Utf1* (Morshedi et al., 2013) or early differentiation genes such as *Gata6* (Heslop et al., 2021). Such reporter lines are useful among others in studying the mechanisms underlying exit from pluripotency and somatic nuclear reprogramming. By producing engineered fibroblasts from these embryonic stem cell (ESC) reporter lines one can easily monitor pluripotency acquisition following the transduction of a set of transcription factors such as OCT4, SOX2, KLF4 and MYC (OSKM) (Buganim et al., 2012; Buganim et al., 2014) or following nuclear transfer (Boiani et al., 2002).

While elimination of one allele of one gene is considered harmless to the cell, a complete KO may be detrimental to the cell as seen in the case of *Oct4* and *Sox2* KO for pluripotent cells (Masui et al., 2007; Nichols et al., 1998). In contrast, a complete elimination of other important pluripotent genes such as *Nano*g, partially maintains the pluripotent state and contributes to chimeras, but shows a dramatic reduced reprogramming efficiency by their fibroblast derivatives that can only be partially overcome by high levels of exogenous OSKM factors (Carter et al., 2014; Schwarz et al., 2014). Although KI/KO of one allele is a wildly used approach to introduce a reporter gene, our previous study suggests that the quality of the reprograming process to pluripotency might be affected by the loss of even one allele as fibroblasts with only one intact allele of *Nanog* generated lower quality iPSCs compared to controls as assessed by the stringent pluripotency test, the tetraploid complementation (4n) assay (Buganim et al., 2014).

During the maturation phase of the reprogramming process, which is thought to be the bottleneck of the process, epigenetic changes happen stochastically to eventually allow expression of the first pluripotent-related genes (Buganim et al., 2013). Using single-cell analyses, it has been shown that stochastic low expression of pluripotent genes such as *Utf1*, *Esrrb*, *Sall4* (Buganim et al., 2012) and *Nanog* (Polo et al., 2012) can be observed early on in the process in a small fraction of induced cells which is correlated to the low efficiency of reprogramming. The stochastic behavior of the maturation phase ends with the activation of late pluripotent genes such as *Sox2*, *Dppa4*, *Prdm14* and *Gdf3* (Buganim et al., 2012; Soufi et al., 2012) which unleashes the final deterministic phase of reprogramming which leads to stabilization by activation of the core pluripotency network, transgene silencing and complete epigenetic resetting (Buganim et al., 2013).

While major efforts have been put to decipher how the identity and levels of the exogenous pluripotent reprogramming factors are linked to efficiency of reprogramming and quality of the resulting iPSCs (Benchetrit et al., 2015; Buganim et al., 2013; Theunissen and Jaenisch, 2014), studies dealing with the effect of reduced levels of endogenous pluripotency genes during development or during pluripotency induction are mostly based on one gene KO approach, that eliminates completely the expression of the targeted gene, or on haploid ESC systems that become diploid very soon during development (Elling et al., 2019; Leeb and Wutz, 2011). Thus, there is very little knowledge of how reduced levels of multiple endogenous pluripotency genes in pluripotent cells affects their developmental potential and their somatic cell derivatives.

Here, we sought to examine how elimination of one allele of two pluripotency genes in different pluripotent systems affects their developmental potential and the efficiency of the reprogramming process of their fibroblast derivatives. We produced three secondary systems where two pluripotency genes were KO only in one of the two alleles. These double heterozygous mutant lines include NGFP2 (Nanog^+/-^;Sall4^+/-^, Nanog^+/-^;Esrrb^+/-^ and Nanog^+/-^;Utf1^+/-^), NGFP1 (Nanog^+/-^;Sall4^+/-^) and SGFP1 (Sox2^+/-^;Sall4^+/-^). Interestingly, while all double heterozygous mutant lines were capable of contributing to chimeras in a comparable manner to their parental secondary iPSC systems (i.e. NGFP2 (Nanog^+/-^), NGFP1 (Nanog^+/-^) and SGFP1 (Sox2^+/-^)), multiple derivations of fibroblasts from these lines resulted in poor reprogramming efficiency ranging from a complete blockage at the mesenchymal to epithelial (MET) transition (NGFP2 line) to a late blockage at the stabilization step just before the acquisition of pluripotency (NGFP1 and SGFP1 lines). This reduced efficiency was not limited to reprogramming by defined factors but also was evident in nuclear transfer (NT). To understand whether reduced early stochastic expression of these key pluripotency genes can explain the low efficiency of the reprogramming process we generated tracing systems for *Sall4* and *Nanog* as major determinants for the reprogramming process. Tracing *Sall4* or *Nanog locus* activation along the reprogramming process revealed that only a very small fraction of cells activated these *loci*, at one point during the stochastic phase, a result that cannot explain the global blockage seen during the reprogramming process with these double heterozygous mutant lines. To further understand this phenomenon, we profiled the CpG-riched methylation landscape of fibroblasts derived from SGFP1 double heterozygous mutant line and their parental control. Interestingly, a clear difference in the methylation levels of multiple developmental and pluripotent *loci* was observed between the double heterozygous mutant fibroblasts and their parental control cells. In agreement with that, treating double heterozygous mutant fibroblasts for two days prior to factor induction with 5-azacytidine rescued the reprogramming blockage and allowed the induction of pluripotency. This study emphasizes the importance of having two intact alleles for proper pluripotency induction and for normal embryonic development and raises a concern regarding the often used approach of reporter introduction using a KI/KO targeting technique.

## RESULTS

### Double heterozygous mutant pluripotent cells contribute to chimeras and exhibit modest transcriptional changes

Given the importance of properly functioning core ESC circuitry for the establishment and maintenance of pluripotency, we hypothesized that even a small reduction in gene expression of few key pluripotency genes might hold a dramatic effect on the developmental potential of the cells or on their derivatives to undergo nuclear reprogramming.

We decided to focus our research on secondary iPSC systems as these systems on the one hand contribute to chimeras and on the other hand exhibit stable and reproducible reprogramming efficiency by minimizing cell heterogeneity (Wernig et al., 2008). Moreover, it allows us to compare a single allele KO of one gene to a single allele KO of two genes with minimal background heterogeneity (Haenebalcke et al., 2013).

We started by targeting the NGFP2 secondary system as it already contains a single KI/KO allele of *Nanog* (Wernig et al., 2008). We chose to eliminate a single allele of *Esrrb*, *Utf1* or *Sall4* as they have all been shown to be important for pluripotency and to play a role in reprogramming during the stochastic phase (Buganim et al., 2012; Feng et al., 2009; Tsubooka et al., 2009). To produce a single allele KO and to be able to monitor the activity of the targeted allele, we designed donor vectors that fused, in frame, to the first or second exon a tdTomato reporter (**Figure 1A-B**). Moreover, although a stop codon at the end of the tdTomato was introduced to the targeted allele, to avoid exon skipping and to completely destabilizing the mRNA of the targeted allele we did not add polyA to the targeting vectors. NGFP2 iPSCs were electroporated with either of the three targeting vectors and treated with neomycin for a week. Stable colonies were isolated, expended and examined for correct targeting by southern blots using external or internal probes (**Figure 1C**, correctly targeted clones are marked by red asterisks). Overall, we isolated two correctly targeted colonies for each combination of manipulated genes: Nanog^+/-^; Esrrb^+/-^ (NGFP2^N+/-;E+/-^), Nanog^+/-^; Utf1^+/-^ (NGFP2^N+/-;U+/-^) and Nanog^+/-^; Sall4^+/-^ (NGFP2^N+/-;S+/-^). To validate the reduced levels of the targeting genes, we cultured the cells in 2i/L medium that recapitulates the ground pluripotent state and facilitates gene expression from both alleles (Miyanari and Torres-Padilla, 2012). qPCR and western blot analyses clearly demonstrated a reduction in about 50% of the total mRNA or protein levels of all targeted alleles (**Figures 1D and S1A**), but not in other key pluripotency genes such as *Oct4*, *Sox2*, *Lin28*, *Fbxo15* and *Fgf4* as assessed by qPCR (**Figure S1A**). It is important however to note that out of the examined genes some further reduction in the protein level of NANOG and ESRRB was seen in NGFP2^N+/-;U+/-^ and NGFP2^N+/-;S+/-^ iPSC lines (**Figure 1D**) and in the mRNA of the *Dppa3* gene in NGFP2^N+/-;S+/-^ line (**Figure S1A**). These results suggest that NANOG and ESRRB are either direct or indirect targets of *Sall4* and *Utf1* and that *Dppa3* is regulated by SALL4. To test the stability of the mRNA of the targeted allele we grew the various double heterozygous mutant lines either in S/Lif or 2i/Lif conditions and subjected them to flow cytometry analysis for GFP and tdTomato activity. As expected, and in agreement with the western blot analysis, cells grown under S/Lif conditions (i.e. conditions that mostly facilitate mono-allelic expression of Nanog) exhibited 68% GFP reporter activity (reporter that was introduced in frame and contains polyA) in NGFP2^N+/-^ control and NGFP2^N+/-;E+/-^ iPSC lines, and 55% and 58% in NGFP2^N+/-;S+/-^ and NGFP2^N+/-;U+/-^ iPSC lines, respectively (**Figure 1E**). In contrast to the Nanog-GFP reporter and in accordance with our strategy, tdTomato activity for all targeted genes was minor due to the absence of a polyA which resulted in the destabilization of the targeted mRNA (**Figure 1E**). A better activation of the Nanog-GFP reporter was noted under 2i/Lif conditions in all clones but a reduced percentage was still evident in all heterozygous mutant iPSC lines (**Figure S1C**). As in the S/Lif conditions, the activation of the tdTomato reporter in 2i/Lif conditions was minor but still showed a stronger activation than S/Lif conditions. These results validate our strategy of eliminating a single allele in several combinations of two pluripotency genes.

**Figure 1.**
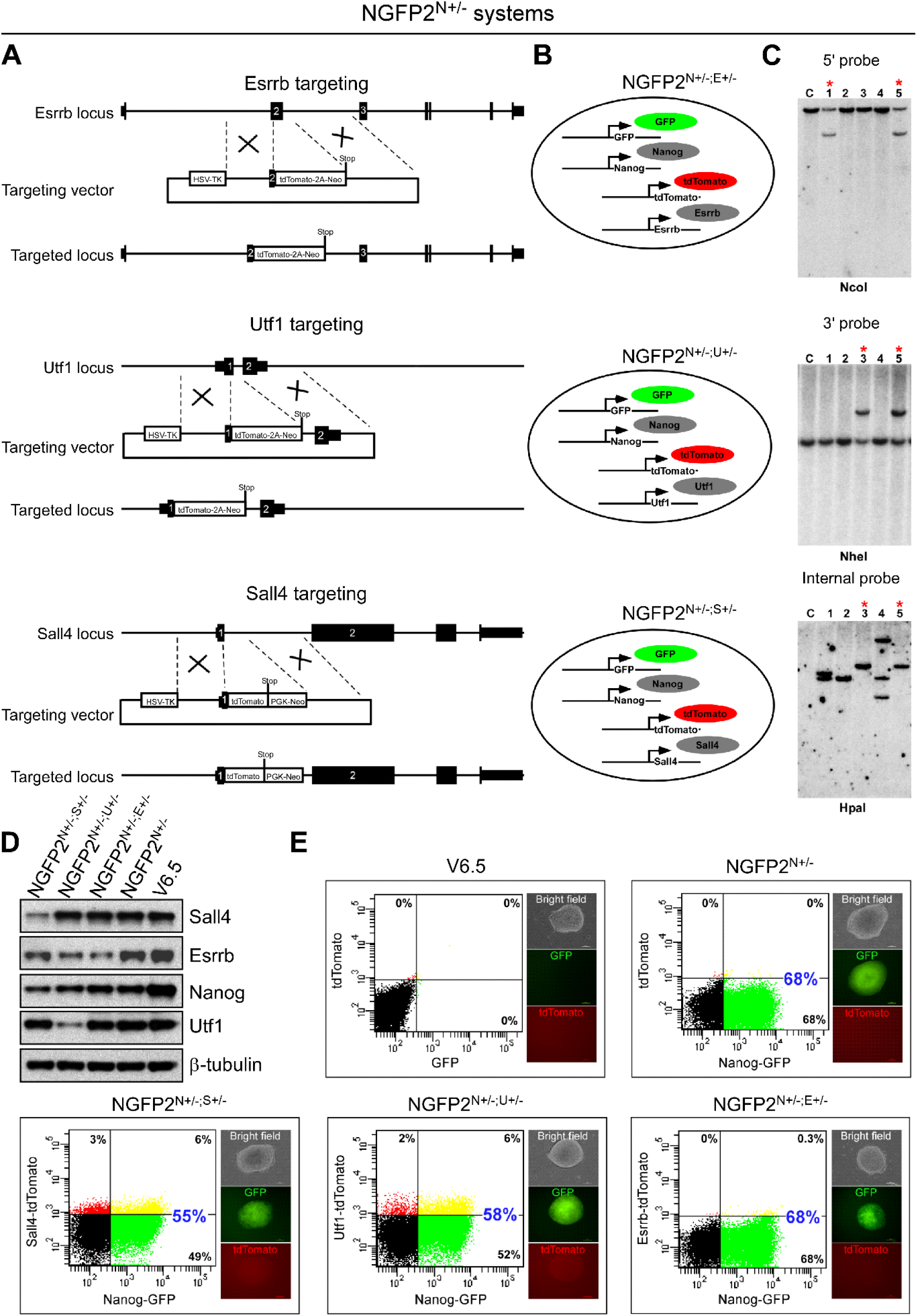
Generation of double heterozygous mutant NGFP2^N+/-^ iPSC lines. **(A, B)** Schematic representation of the KI/KO targeting strategy for replacing one allele of *Esrrb, Utf1* or *Sall4* with a tdTomato reporter in NGFP2^N+/-^ line. **(C)** Southern blot analysis for NGFP2^N+/-^ targeted iPSC clones demonstrating heterozygous targeting for *Esrrb, Utf1* and *Sall4*. Correctly targeted clones are marked by red asterisks. **(D)** Western blot analysis demonstrating a reduction of ∼50% of the protein levels of the targeted gene (*Esrrb, Utf1, Nanog* and *Sall4*) compared to ESC (V6.5) control. Cells were grown in 2i condition to facilitate expression from both alleles. **(E)** Flow cytometry analysis for GFP (*Nanog*) and tdTomato (*Utf1, Esrrb* or *Sall4*) in the various double heterozygous mutant lines that grew under S/Lif conditions. Note that while cells express the GFP reporter due to functional polyA signal, tdTomato is hardly detectable due to the absence of polyA. Similar to the western blot (D) GFP expression is reduced even further in NGFP2^N+/-;U+/-^ and NGFP2^N+/-;S+/-^ compared to NGFP2^N+/-;E+/-^ and the parental cells NGFP2^N+/-^.

We first wished to investigate the impact of eliminating a single allele in two different pluripotent genes on the developmental potential of the cells. To that end, we injected the three double heterozygous mutant lines as well as their parental control cells into blastocysts and measured their potential to form chimeric mice. As can be seen in representative images in Figure S2A, a comparable grade of chimerism was noted between all double heterozygous mutant lines and control iPSC line, suggesting that elimination of a single allele in these combinations of two pluripotent genes does not exert a significant developmental barrier (**Figure S2A**).

In our previous study we showed that a gene list of 1716 genes can distinguish between iPSCs with poor, low and high quality as assessed by grade of chimerism and 4n complementation assay (Buganim et al., 2014). Thus, we next profiled the transcriptome of the three heterozygous mutant lines, the parental NGFP2^N+/-^ cells and wild type (WT) ESC (V6.5) control line under S/Lif or 2i/Lif conditions using RNA-sequencing (RNA-seq). Correlation heatmap clustered the cells into two main groups based on the culture conditions used. Nevertheless, within the S/Lif group some changes in gene expression were noted in NGFP2^N+/-;S+/-^ and NGFP2^N+/-;U+/-^ compared to NGFP2^N+/-;E+/-^, parental NGFP2^N+/-^ and control ESC line (**Figure S2B**). As *Esrrb* was shown to be a downstream target gene of NANOG and to exert a positive feedback loop with it (Festuccia et al., 2012; Sevilla et al., 2021), it is not surprising that the parental NGFP2^N+/-^ and NGFP2^N+/-;E+/-^ exhibited minimal transcriptional changes between them and clustered very close to each other. Principal component analysis (PCA) validated the results seen in the correlation heatmap, separating S/Lif conditions from 2i/Lif conditions by PC1 and NGFP2^N+/-;S+/-^ and NGFP2^N+/-;U+/-^ that grown under S/Lif condition from the rest of the samples by PC2 (**Figure S2C**). Interestingly, NGFP2^N+/-;U+/-^ grown under S/Lif conditions, clustered closer to samples that grew under 2i conditions as indicated by PC1 (**Figure S2C**). These results are in accordance with the notion that UTF1 is mostly implicated in a more primed pluripotent state and less in the ground state (Martinez-Val et al., 2021). As expected, cells grown under 2i/Lif conditions clustered together with minimal transcriptional changes between them (**Figure S2C**). These results suggest that elimination of one allele of two different pluripotent genes from the tested combinations, although show some small transcriptional change under S/Lif conditions, can still maintain a functional pluripotency state with minimal variation in gene expression under ground pluripotency state.

### Fibroblasts derived from NGFP2 double heterozygous mutant iPSC lines fail to induce pluripotency

Given that the reprogramming process involves a stochastic phase of activation of pluripotency genes (Buganim et al., 2012), we hypothesized that MEFs harboring double heterozygous mutant alleles might exhibit reprogramming delay due to difficulties in the activation of the core pluripotency circuitry.

To that end, secondary MEF systems were established from all the three NGFP2 double heterozygous mutant lines, as well as from the parental NGFP2 control. These secondary MEF systems contain a unique integration pattern of OSKM transgenes under Tet-on promotor and a M2rtTA transactivator in the *Rosa26 locus* (Wernig et al., 2008). To initiate reprogramming, MEFs were exposed to dox for 13 days followed by dox withdrawal for 3 more days to stabilize any iPSC colony, and the percentage of Nanog-GFP-positive cells was scored by flow cytometry.

In accordance with our hypothesis, while NGFP2^N+/-^ control cells exhibited the expected ∼2% of Nanog-GFP-positive cells (Buganim et al., 2012; Wernig et al., 2008), all the double heterozygous mutant lines showed a blockage in reprogramming (**Figure 2A**). This blockage was not due to cell death or proliferation arrest as all double heterozygous mutant and control plates stained equally to crystal violet, indicating a comparable number of cells between all mutant lines and control (**Figure 2B**). However, in agreement with the flow cytometry results, although all the double heterozygous mutant lines stained positive to the early reprogramming marker alkaline phosphatase (AP), implying that the cells initiated reprogramming, their AP staining was significantly lower compared to the control plates (**Figure 2C**). By extending the dox exposure time to 20 days, a small percentage of Nanog-GFP-positive cells (i.e. ∼0.3-0.4% Nanog-GFP-positive cells in the mutant lines compared to ∼10% in the control line) did emerge in all double heterozygous mutant lines, suggesting that some cells can overcome this blockage when prolonged exposure of the 4 factors is triggered (**Figure 2D**).

**Figure 2.**
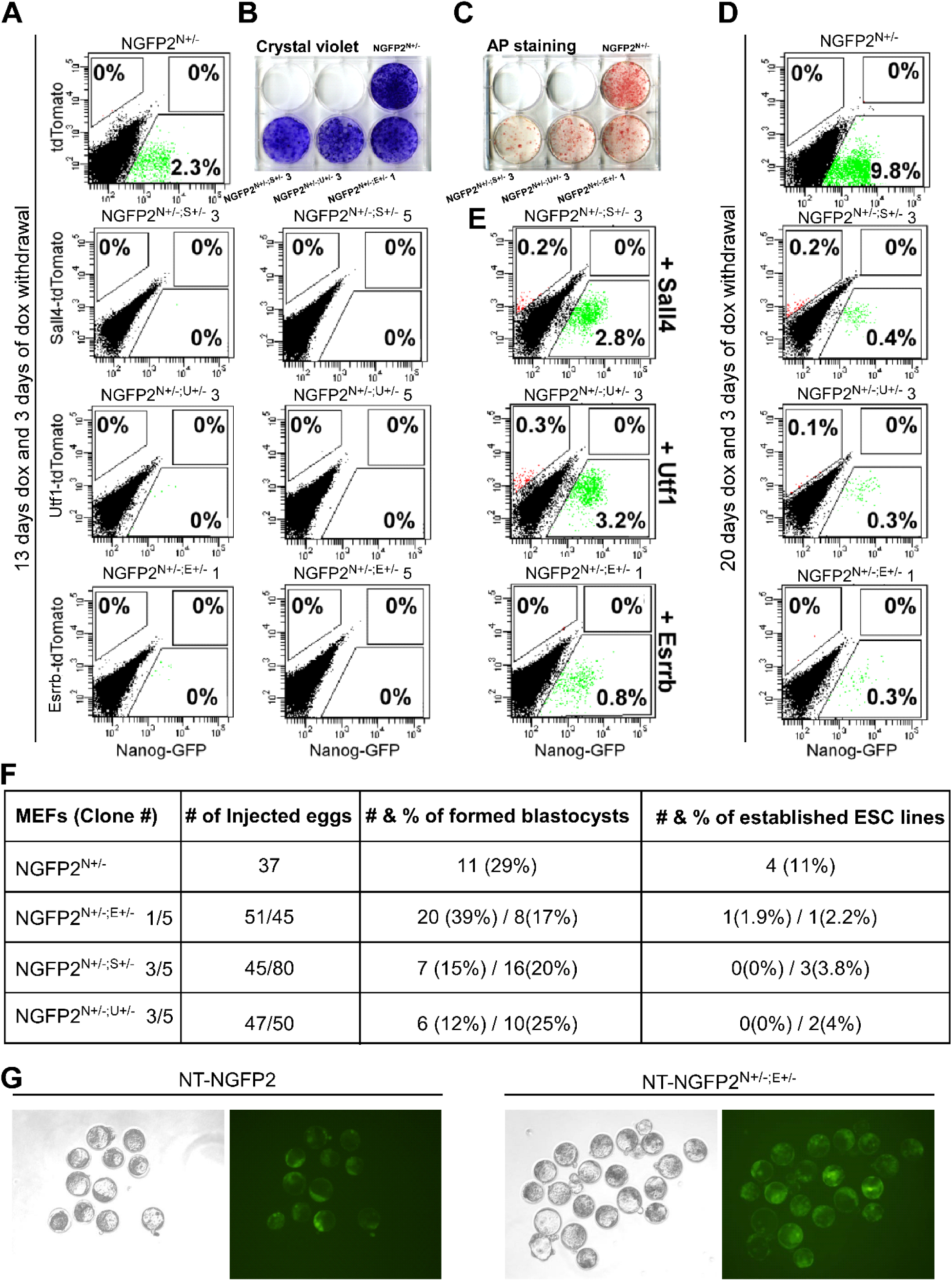
NGFP2^N+/-^ double heterozygous mutant MEFs show strong reprogramming inhibition either by OSKM or by somatic cell nuclear transfer (SCNT). **(A)** Flow cytometry analysis of Nanog-GFP and tdTomato-positive cells for two different clones from each of the NGFP2 double heterozygous mutant MEFs and control following 13 days of dox exposure following by 3 days of dox withdrawal. **(B)** Crystal violet staining of whole reprogramming plate for each of the double heterozygous mutant MEF line and control at the end of the reprogramming process **(C)** Alkaline phosphatase staining of whole reprogramming plate of each of the double heterozygous mutant MEF line and control at the end of the reprogramming process. **(D)** Flow cytometry analysis of Nanog-GFP and tdTomato-positive cells for each of the NGFP2^N+/-^ double heterozygous mutant MEFs and control following 20 days of dox exposure following by 3 days of dox withdrawal. **(E)** Flow cytometry analysis of Nanog-GFP and tdTomato-positive cells of each of the NGFP2^N+/-^ double heterozygous mutant MEFs and control following overexpression of the targeted gene (*Sall4, Utf1* and *Esrrb*) and reprogramming process of 13 days and 3 days of dox withdrawal. **(F)** Table summarizing the efficiency (i.e. blastocyst formation and ESC derivation) of the somatic cell nuclear transfer (SCNT) process of MEF nuclei of the different double heterozygous mutant NGFP2^N+/-^ lines. **(G)** Representative bright field and green channel images of NGFP2^N+/-^ and NGFP2^N+/-;E+/-^ following SCNT. Note that both cell lines produced Nanog-GFP-positive blastocysts.

We then asked whether the observed reprogramming blockage within these double heterozygous mutant MEF lines is specific to reprogramming by defined factors or whether the loss of the two alleles would show reprogramming defects in other reprogramming techniques as well. We chose to use the nuclear transfer (NT) technique as it utilizes the entire array of reprogramming proteins within the egg as opposed to the Yamanaka’s approach that uses very few selected reprogramming factors. Enucleated eggs were injected with MEF nuclei from each of the three double heterozygous mutant MEF lines and control, and blastocyst formation and the establishment of ESC lines from these blastocysts were scored. Notably, while all lines exhibited a comparable and expected efficiency in producing blastocysts, the efficiency of ESC line derivation was significantly lower in the double heterozygous mutant lines compared to control (i.e. 0-4% in the double heterozygous mutant lines vs 11% in control line, **Figure 2F**). Taken together, these results suggest that the elimination of two alleles from two different key pluripotency genes affects the somatic nucleus in a way that interferes with its capability to undergo reprogramming to pluripotency by various techniques.

We then asked whether the loss of the two alleles causes a permanent reprogramming defect or whether it can be rescued by exogenously expressing one of the targeted genes. To that end, each double heterozygous mutant MEF line was transduced with either *Nanog* or with its corresponding targeted gene (i.e. *Sall4*, *Utf1* or *Esrrb*) or with additional viruses encoding for OSK. Overall, both *Nanog* or each of the corresponding factor showed either partial (i.e. Esrrb in NGFP2^N+/-;E+/-^ cells and Nanog in NGFP2^N+/-;S+/-^ cells) or complete rescue of the reprogramming blockage, while additional OSK further boosted the reprogramming process (**Figures 2E and S2D-E**). The fact all the double heterozygous mutant lines could be rescued by the addition of different pluripotent factors, suggests that the seen blockage is not specific to the unique function of the eliminated alleles’ but rather is associated with a more general effect.

### NGFP2^N+/-^ double heterozygous mutant lines show an early defect in the activation of epithelial markers

Given that double heterozygous mutant MEF lines are capable, to some extent, to activate the AP enzyme (**Figure 2C**), we next sought to understand at which time point during the reprogramming process the double heterozygous mutant cells lose their capacity to undergo reprogramming. To that end, we examined the expression level of early reprograming markers (*Fgf4* and *Fbxo15*), intermediate markers (*endogenous Oct4* and *Sall4*) and late and predictive markers (*Sox2*, *Utf1*, *Esrrb* and *Lin28*) following 13 days of dox addition (Buganim et al., 2012; Buganim et al., 2013)). While the double heterozygous mutant induced cells showed some activation of the early markers and very low expression of intermediate markers, no activation at all was seen in the late and predictive markers compared to control cells (**Figure S2F**). These results suggest that the blockage seen in these double heterozygous mutant cells during reprogramming occurs relatively early in the reprogramming process. It is interesting to note that out of the three double heterozygous mutant lines, NGFP2^N+/-;S+/-^ showed the strongest inhibitory effect as assessed by marker expression (**Figure S2F**), Nanog-GFP activation in the *Nanog* rescue experiment (**Figure S2D**), and AP staining (**Figure 2C**).

We next profiled the transcriptome of the three double heterozygous mutant lines and control lines (i.e. NGFP2^N+/-^ cells, and NGFP2^N+/-^ cells that were infected with empty vector (EV)) following six days of reprogramming. We chose this time point as it showed a clear reprogramming delay in the double heterozygous mutant plates compared to control plates. NGFP2^N+/-^ MEFs and the parental NGFP2^N+/-^ iPSCs were profiled as well. Hierarchical clustering analysis showed that all the double heterozygous mutant lines clustered together and different than the control lines (**Figure 3A**). PCA and scatter plots emphasize even more the transcriptomic differences between the double heterozygous mutant lines and NGFP2^N+/-^ control lines after 6 days of dox. While the control lines demonstrated significant transcriptional changes by day 6 of reprogramming (represented by PC2) compared to parental NGFP2^N+/-^ MEFs, all the double heterozygous mutant lines showed minimal transcriptional changes between themselves or when compared to NGFP2^N+/-^ MEFs (**Figure 3B-D**). These results suggest that an early reprogramming defect is responsible for the delay seen in the NGFP2^N+/-^double heterozygous mutant lines.

**Figure 3.**
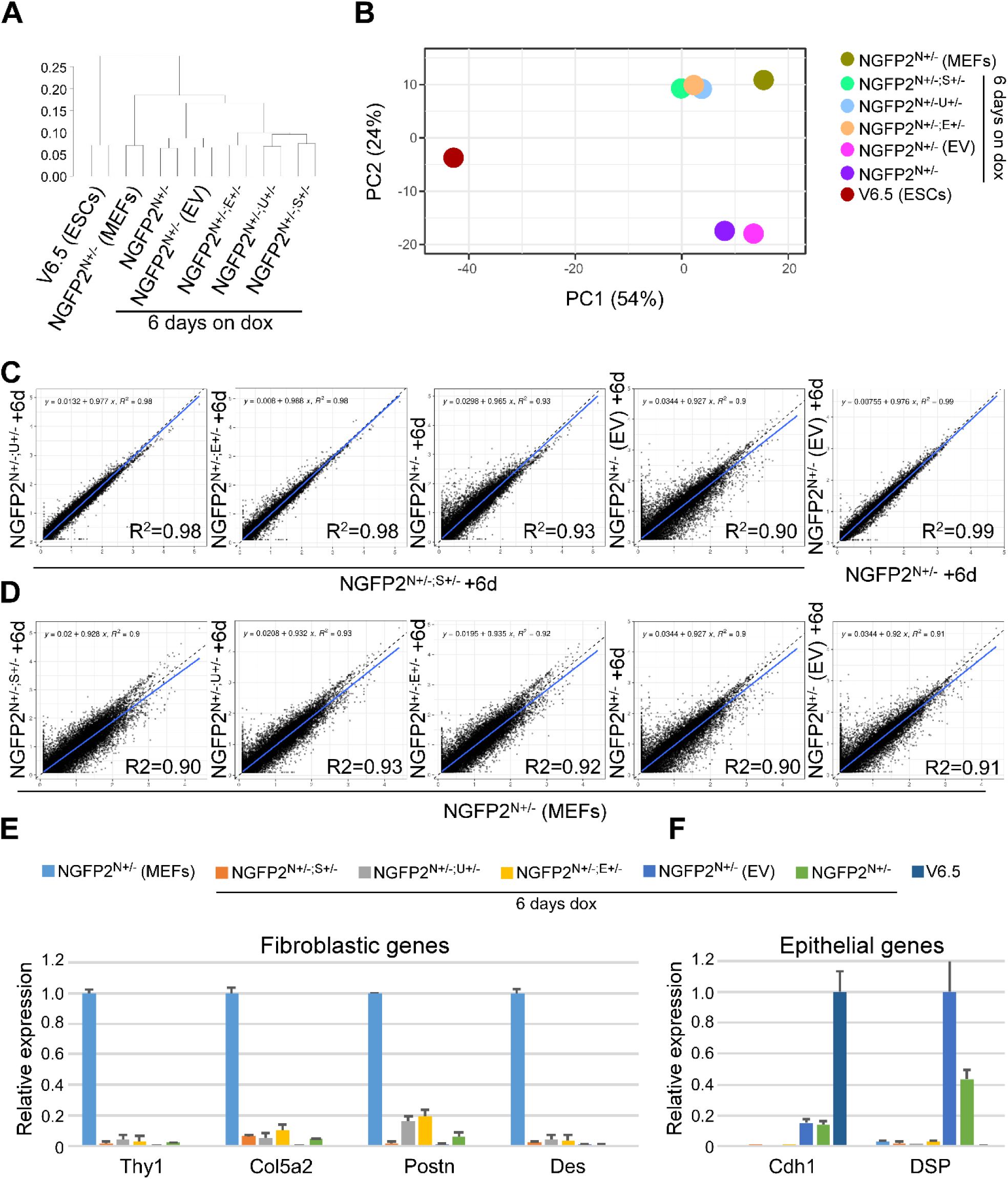
Unbiased comparative transcriptome analyses after 6 Days of dox clusters NGFP2^N+/-^ double heterozygote lines far from NGFP2^N+/-^ control. **(A)** Hierarchical clustering of global gene expression profiles for two RNA-seq replicates for NGFP2^N+/-^ iPSCs, NGFP2^N+/-^ MEFs and NGFP2^N+/-^, NGFP2^N+/-^ (EV) and the various NGFP2^N+/-^ double heterozygous mutant lines (NGFP2^N+/-; E+/-^, NGFP2^N+/-; U+/-^ and NGFP2^N+/-; S4+/-^) after 6 days of reprogramming. Replicate pairs were assigned a shared numerical value. **(B)** Principle component analysis for genes from (A). PC1, 54%; PC2, 24%. Each line is marked by a specific color. The group names correspond to the names in (A). **(C, D)** Scatter plots comparing gene expression between the indicated NGFP2^N+/-^ lines after 6 days of dox and controls. Blue line shows the linear representation of the data, black line shows the y = x line. **(E, F)** qPCR of the indicated fibroblastic genes (E) and epithelial genes (F) in NGFP2^N+/-^ and the different NGFP2^N+/-^ double heterozygous mutant lines after 6 days of dox, MEFs and V6.5 ESCs controls. mRNA levels were normalized to the housekeeping control gene *Gapdh*. Error bars presented as a mean ± SD of 2 duplicate runs from a typical experiment out of 3 independent experiments.

An important and essential early step in reprogramming is mesenchymal to epithelial transition (MET). In this step the induced cells lose their fibroblastic characteristics and start acquiring an epithelial identity. As the transcriptome of the double heterozygous mutant lines after 6 days of reprogramming (**Figure 3B-D**) was still very close to MEFs, we next asked whether the MET process is impaired in the double heterozygous mutant lines. Initially we examined the expression levels of four well-known fibroblastic markers; *Thy1*, *Col5a1*, *Postn*, and *Des* and noticed that in all mutant lines these markers are significantly downregulated in a comparable manner to that of the control lines (**Figure 3E**). Similarly, a comparable downregulation in the expression of the EMT master regulators; *Twist1*, *Zeb1*, *Snai2*, and *Foxc2* was noted as well (**Figure S3A**), implying that the loss of the fibroblastic identity is not impaired in the double heterozygous mutant lines. However, when we tested whether the cells are capable of activating the epithelial program, we noticed that while the control lines are able to activate epithelial markers such as *Cdh1*, *Dsp*, *Epcam*, *Cldn4*, and Cldn7, the double heterozygous mutant lines fail in doing so (**Figure 3F and S3B**). These results suggest that there is an inherent blockage acquired specifically in the double heterozygous mutant MEF lines that prevents/delays them from activating a robust epithelial program as occurs during intact cellular reprogramming.

### Reprogramming impairment caused by double heterozygous allele elimination is not restricted to a system nor to the identity of the modified alleles

To exclude the possibility that the observed effect is system-specific, we produced additional secondary double heterozygous mutant iPSC systems that differ in their reprogramming efficiency and dynamics, developmental potential, allele-specific elimination and reprogramming factor stoichiometry.

We decided to produce a double heterozygous mutant line from NGFP1^N+/-^ system as it was generated in parallel to the NGFP2^N+/-^ system but demonstrated different reprogramming efficiency and dynamics and reprograming factor induction levels (Wernig et al., 2008).

As NGFP2^N+/-;S+/-^ double heterozygous mutant line demonstrated the strongest delay in pluripotency induction, we decided to eliminate one allele of Sall4 in NGFP1^N+/-^ as well. Initially, we confirmed by single molecule mRNA-FISH that the strong effect seen in NGFP2^N+/-;S+/-^ is not a result of *Sall4* reduction that is greater than 50%. In agreement with the protein level (**Figure 1D**), Figure 4A shows the distribution of *Sall4* transcript level in NGFP2^N+/-^ cells (n=57) compared to NGFP2^N+/-;S+/-^ cells (n=49), validating transcript reduction of *Sall4* in about 50% within NGFP2^N+/-;S+/-^ cells (**Figure 4A**).

**Figure 4.**
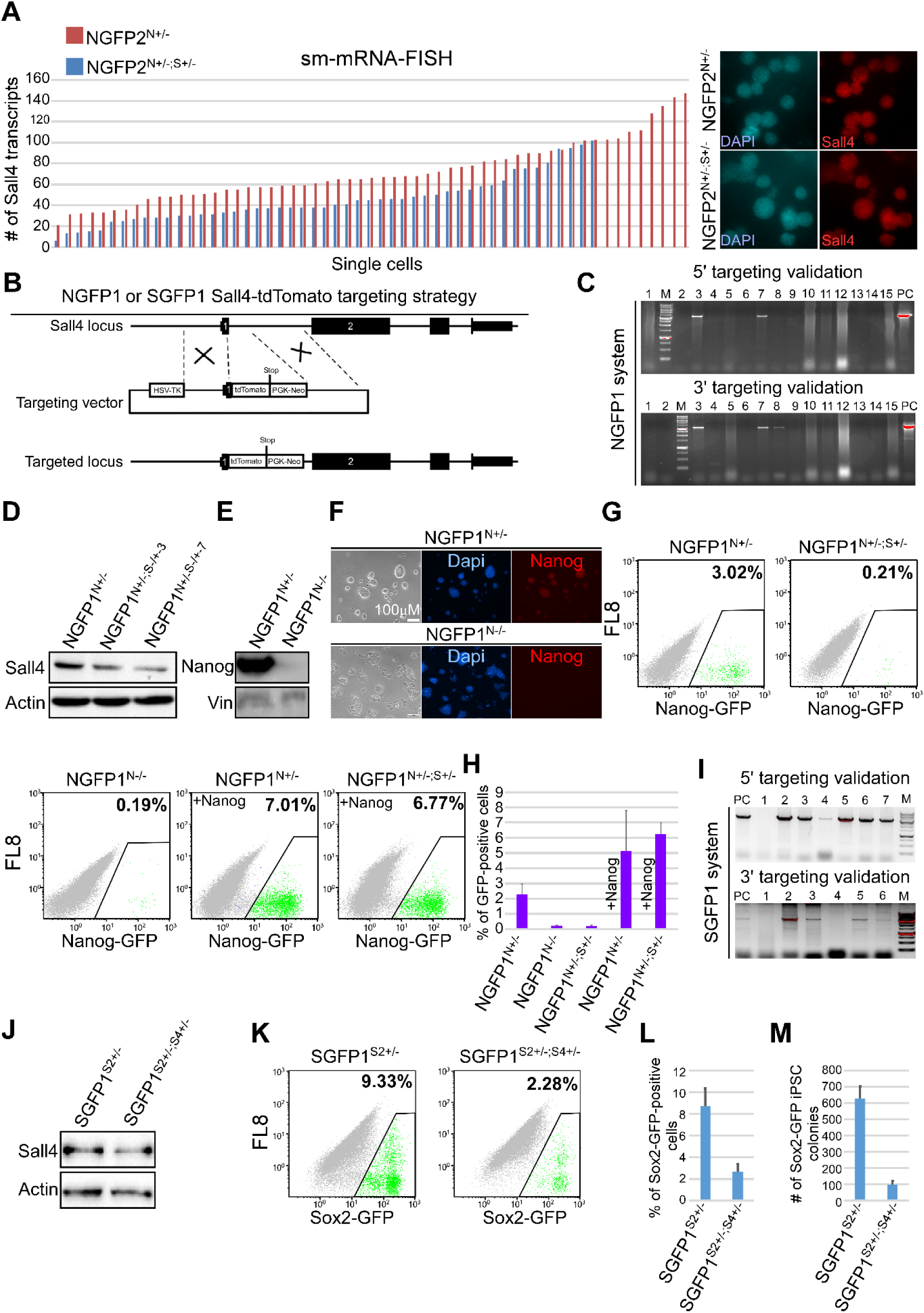
NGFP1^N+/-^ double heterozygous mutant MEFs and Nanog KO MEFs show strong reprogramming inhibition. **(A)** Sm-mRNA-FISH directed towards Sall4 transcripts in NGFP2^N+/-^ and NGFP2^N+/-; S+/-^ single iPSCs. **(B)** Schematic representation of the KI/KO targeting strategy for replacing one allele of *Sall4* with a tdTomato reporter in NGFP1^N+/-^ and SGFP1^S2+/-; S4+/-^ respectively. **(C)** PCR analysis for NGFP1^N+/-^ targeted iPSC clones demonstrating correctly targeted clones for one allele of *Sall4*. Correctly targeted clones were verified using primers amplifying regions at the 5’ and 3’ end of the targeted *locus*. **(D)** Western blot analysis demonstrating a reduction of ∼50% of the protein levels of Sall4 compared to parental NGFP1^N+/-^ control. **(E, F)** NGFP1^N+/-^ parental iPSCs were transfected with CRISPR/Cas9 and gRNA against *Nanog* to produce *Nanog* KO NGFP1^N-/-^ line. Western blot analysis (E) and immunostaining (F) demonstrating a complete KO of *Nanog* compared to parental NGFP1^N+/-^ control. **(G)** Flow cytometry analysis of Nanog-GFP-positive cells in NGFP1^N+/-^ control cells, NGFP1^N+/-;S+/-^, NGFP1^N-/-^ and following overexpression of Nanog in rescue experiments after 13 days of reprogramming following by 3 days of dox withdrawal. **(H)** Comparative percentage of Nanog-GFP-positive cells for NGFP1^N+/-^, NGFP1^N+/-; S+/-^, NGFP1^N-/-^, NGFP1^N+/-^ (+*Nanog* OE) and NGFP1^N+/-; S4+/-^ (+Nanog OE) following reprogramming with OSKM (13 days of dox + 3 days dox withdrawal). **(I)** PCR validation for SGFP1^S2+/-; S4+/-^ clones. **(J)** Western blot analysis detecting Sall4 in SGFP1^S2+/-^ and SGFP1^S2+/-; S4+/-^ iPSCs. **(K)** Flow cytometry analysis of Sox2-GFP-positive cells for SGFP1^S2+/-^ compared with SGFP1^S2+/-; S4+/-^ following reprogramming with OSKM (13 days of dox + 3 days dox withdrawal). **(L)** Comparative percentage of SOX2-GFP-positive cells for SGFP1^S2+/-^ compared with SGFP1^S2+/-; S4+/-^ following reprogramming with OSKM (13 days of dox + 3 days dox withdrawal). **(M)** Graph summarizing the number of colonies counted at the end of the reprograming for SGFP1^S2+/-^ and SGFP1^S2+/-; S4+/-^.

Then, we targeted a tdTomato reporter gene into the *Sall4 locus* of NGFP1^N+/-^ as described above to produce NGFP2^N+/-;S+/-^ (**Figure 4B**). Correctly targeted NGFP1^N+/-;S+/-^ iPSC double heterozygous colonies were validated by PCR and Western blot **(Figure 4C-D)**. We also produced a *Nanog* KO NGFP1^N-/-^ line as a single KO gene control (**Figures 4E-F and S4A**). Secondary MEFs were produced from NGFP1^N+/-^, NGFP1^N+/-;S+/-^, and NGFP1^N-/-^ which were then exposed to dox for 13 days following by dox withdrawal for 3 more days. Flow cytometry analysis of the various reprogramming plates showed a clear and comparable reduction in the percentage of Nanog-GFP-positive cells in the double heterozygous mutant cells and in the *Nanog* KO fibroblasts compared to control NGFP1^N+/-^ cells (**Figure 4G**). Exogenous expression of *Nanog*, from the onset of the reprogramming process, rescued both *Nanog* KO cells and the NGFP1^N+/-;S+/-^ double heterozygous mutant cells (**Figure 4G-H**). These intriguing results suggest that even a reduction of 50% in the levels of two important pluripotency genes (i.e. Nanog and Sall4) is crucial for the establishment of the core pluripotency circuitry in a way that is comparable to a complete KO of key pluripotency genes such as *Nanog*.

We were interested to examine whether the pluripotency induction impairment seen in the double heterozygous mutant lines is restricted to combinations that harbor allele elimination of *Nanog*.

To that end, we eliminated one allele of *Sall4* in SGFP1^S2+/-^ line, a secondary iPSC system that was generated in our lab and contains GFP reporter instead of one allele of *Sox2.* Correctly targeted SGFP1^S2+/-;S4+/-^ iPSC colonies were validated by PCR, Western blot and immunostaining **(Figures 4I-K and S4B).** In agreement with the other systems, a significant reduction in reprogramming efficiency was noted in SGFP1^S2+/-;S4+/-^ cells compared to SGFP1^S2+/-^ control as assessed by flow cytometry and number of Sox2-GFP-positive colonies **(Figure 4K-M)**. It is interesting to note that while all the double heterozygous NGFP^N+/-^ lines produced a neglectable number of iPSCs following 13 days of reprogramming (i.e. 0.0%-0.2%), the SGFP1^S2+/-;S4+/-^ double heterozygous mutant cells produced about 2%-2.5% of iPSCs. This difference can be explained by the levels of the OSKM transgenes that is much higher in SGFP1^S2+/-^ cells than in NGFP^N+/-^ cells (**Figure S4C**) as additional levels of OSK can rescue the phenotype of the double heterozygous mutant lines (**Figure S2E**). Taken together, these results suggest that the double heterozygous phenotype is not system nor gene- specific. This observation raises a real concern as to the KI/KO targeting approach when cellular state induction is studied.

### Reduced early stochastic expression of the targeted genes cannot explain the reprogramming blockage seen in the double heterozygous mutant lines

Stochastic expression of pluripotency genes during early stages of reprogramming was evident by multiple single-cell studies (Buganim et al., 2012; Buganim et al., 2013; Guo et al., 2019). Thus, we hypothesized that the lack of two key pluripotency alleles in the double heterozygous mutant cells might impair their ability to successfully pass the early stochastic phase, resulting in a blockage/delay in reprogramming. To explore this possibility, we generated tracing system for *Nanog* and *Sall4* as two representative genes out of the 5 targeted ones (i.e. *Nanog*, *Sall4*, *Utf1*, *Esrrb* and *Sox2*). We chose *Nanog* because it appears in most of our double heterozygous mutant lines and *Sall4* because it exhibits the highest levels of stochastic expression at early phases of reprogramming compared to the other targeted genes (Buganim et al., 2012).

To that end, we targeted a *2A-EGFP-ERT-CRE-ERT* cassette into *Sall4* or *Nanog locus* using gRNA that is located at the 3’ UTR of the gene. We targeted an ESC line (RL) that contains a lox-STOP-lox (L-S-L) cassette upstream to a tdTomato reporter gene and M2rtTA transactivator at the *Rosa26 locus* (each cassette at different allele, **Figure 5A, 5B**). Upon *Sall4* or *Nanog* expression and the addition of tamoxifen, CRE recombinase is translocated to the nucleus and removes the L-S-L cassette, leading to irreversible activation of the tdTomato reporter. Given that both *Nanog* and *Sall4* are expressed in ESCs, transfected colonies were sorted based on EGFP expression and correct targeting was validated by PCR (**Figure 5C-D**). To assess the efficiency of the tracing system, correctly targeted ESC clones (i.e. RL8 for *Sall4* and RL9 for *Nanog*) were exposed to Tamoxifen (Tam) and the percentage of tdTomato-positive cells were scored under the microscope and by flow cytometry, demonstrating very high L-S-L cassette removal efficiency (**Figures 5E-F and S5A-D**).

**Figure 5.**
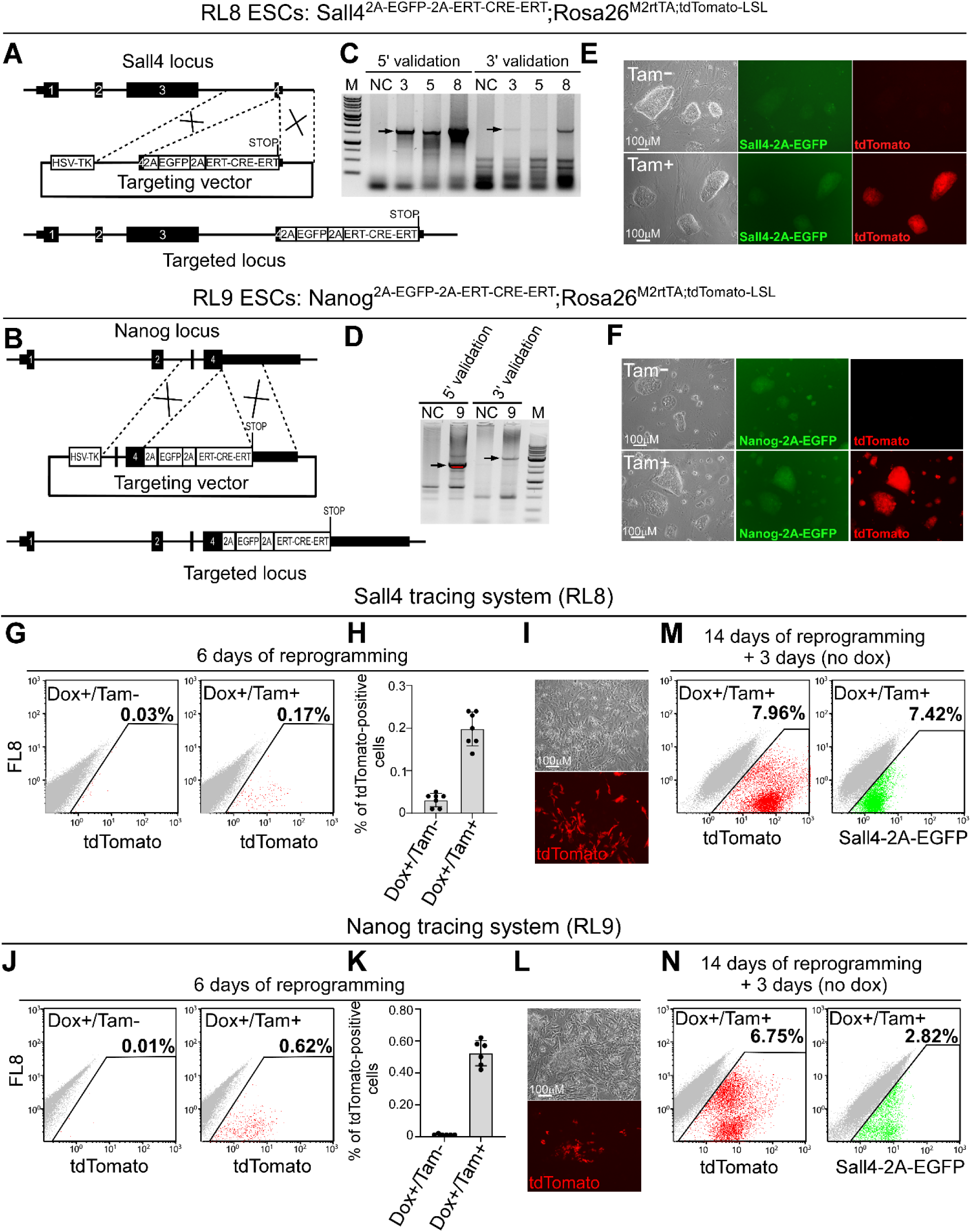
Sall4 and Nanog tracing systems cannot explain the reprogramming blockage observed in the double heterozygous mutant cells. **(A, B)** Schematic representation of the targeting strategy to introduce a 2A-EGFP-ERT-CRE-ERT cassette into the *Sall4 locus* (A) or into the *Nanog locus* (B). **(C, D)** PCR validation for targeted colonies demonstrating a correct targeting band size for the *Sall4 locus* (C) and for the *Nanog locus* (D) using both 5’ and 3’ regions of the incorporation point. Black arrows depict the band size of correctly targeted allele. NC- negative control. **(E, F)** Representative bright field, RFP and GFP channel images for the Sall4 tracing system (RL8, E) and for the Nanog tracing system (RL9, F) before and after tamoxifen addition. Scale bar, 100μm. **(G)** Flow cytometry analysis of tdTomato-positive RL8 MEFs that were infected with OSKM and submitted to dox for 6 days with or without tamoxifen (Tam). **(H)** Graph summarizing the percentages of tdTomato-positive cells of the Sall4 tracing system after 6 days of dox with or without Tamoxifen. **(I)** Bright field and RFP channel images of tdTomato-positive cells from the Sall4 tracing system after six days of dox and tamoxifen addition. **(J)** Flow cytometry analysis of tdTomato-positive RL9 MEFs that were infected with OSKM and submitted to dox for 6 days with or without tamoxifen (Tam). **(K)** Graph summarizing the percentages of tdTomato-positive cells of the Nanog tracing system after 6 days of dox with or without Tamoxifen. **(L)** Bright field and RFP channel images of tdTomato-positive cells from the Nanog tracing system after six days of dox and tamoxifen addition. **(M, N)** Flow cytometry analysis of tdTomato and Sall4-GFP-positive cells (M) or Nanog-GFP-positive cells (N) following 14 days of OSKM induction in the presence of dox and Tamoxifen followed by 3 days of dox withdrawal.

The observation that NGFP2^N+/-^ double heterozygous mutant induced cells could not activate the epithelial program (**Figure 3F** and **S3B**) suggests that the blockage in reprogramming in these cells is general and not restricted to the small fraction of cells that are destined to become iPSCs. Thus, in order to correlate the stochastic expression of the targeted alleles to the observed blockage/delay, most induced cells should show some activation of the targeted alleles at early time point of reprogramming.

MEFs produced from *Sall4* and *Nanog* tracing ESC systems were transduced with OSKM cassette and tdTomato activation was assessed in the induced cells after 6 days and following 14 days of reprogramming followed by 3 days of dox removal. We chose day 6 since it is an early time point that exhibits high stochastic expression of pluripotency genes (Buganim et al., 2012; Buganim et al., 2013; Guo et al., 2019). However, only up to 0.24% of the Sall4 tracing cells and up to 0.62% of Nanog tracing cells were tdTomato-positive at day 6 of reprogramming, ruling out the hypothesis that *Sall4* or *Nanog* stochastic expression early in the reprogramming process is responsible for the double heterozygous mutant phenotype (**Figure 5G-I, 5J-L**). 7.42% of Sall4-EGFP in conjunction with 7.96% of tdTomato-positive cells for the Sall4 tracing system and 2.8% of Nanog-EGFP together with 6.7% of tdTomato-positive cells for the Nanog tracing system at the end of the reprogramming process confirmed successful reprogramming, refuting the possibility that the low percentage of tdTomato-positive cells observed at day 6 of reprogramming in the *Sall4* and *Nanog* tracing systems is due to low reprogramming efficiency (**Figure 5M-N**). In conclusion, this set of experiments, challenges the hypothesis that reduced stochastic expression of the targeted pluripotent alleles is responsible for the early blockage seen in the double heterozygous mutant lines.

### Methylation abnormalities in the double heterozygous mutant fibroblasts is correlated with reprogramming impairment

The fact that additional exogenous expression of OSK factors could rescue the delay seen in the double heterozygous mutant cells (**Figure S2E**), suggests that epigenetic abnormalities, rather than genetic modifications (i.e. the elimination of the targeted alleles themselves), are responsible for the observed blockage. This notion is also supported by the transcriptomic changes seen in the double heterozygous NGFP2^N+/-^ iPSC mutant lines that grew under S/Lif condition, but not in 2i/Lif conditions, that force the naïve ground state (**Figure S2B-C**). As 2i/Lif conditions induce DNA hypomethylation (Sim et al., 2017), and since DNA methylation reshaping has been shown to be crucial for reprogramming (Buganim et al., 2013; De Carvalho et al., 2010) we hypothesized that the double heterozygous mutant fibroblasts might harbor abnormal DNA methylation that hinders their ability to undergo reprogramming. To test this hypothesis, MEFs derived from double heterozygous mutant SGFP1^S2+/-;S4+/-^ iPSCs and from their parental SGFP1^S2+/-^ iPSCs were subjected to methylation analyses using reduced representation bisulfite sequencing (RRBS) to capture the CpG-enriched methylation landscape of the cells as a representation for the global methylation changes.

RRBS analysis revealed that the two fibroblast lines are very similar in regard to their CpG-enriched methylation landscape, suggesting that overall the double heterozygous mutant line harbor a correct fibroblastic methylation landscape. However, read counts did vary between samples and so did reads-per-site, clustering them as two different groups (**Figure 6A**). We then searched for differentially methylated regions (DMRs) between the two fibroblast lines. DMRs were defined as CpG sites of consecutive tiles that are 100bp long in size, include at least 15 reads and show at least 20% methylation differences between the two fibroblast lines. All DMRs were adjusted to p-value of 1e-3 or lower. This analysis yielded two groups of DMRs: (i) 1263 tiles that are more methylated and (ii) 1384 tiles that are less methylated in the double heterozygous mutant line compared to control (**Figure 6B-C**). We then associated each DMR to its neighboring gene and ran GO term analysis using EnrichR (Xie et al., 2021). Interestingly, many of the differentially methylated *loci* were found to be associated with pluripotency and developmental pathways (**Figure 6D-E**). Specifically, dataset derived from loss of function experiments suggested that these genes are being upregulated in ESCs upon loss of function of *Oct4*, and are associated with the Hippo signaling (**Figure 6D-E**), suggesting that the loss of indicated two pluripotency alleles hinders the capability of the core pluripotency circuitry to maintain normal DNA methylation of these *loci* later on in development.

**Figure 6.**
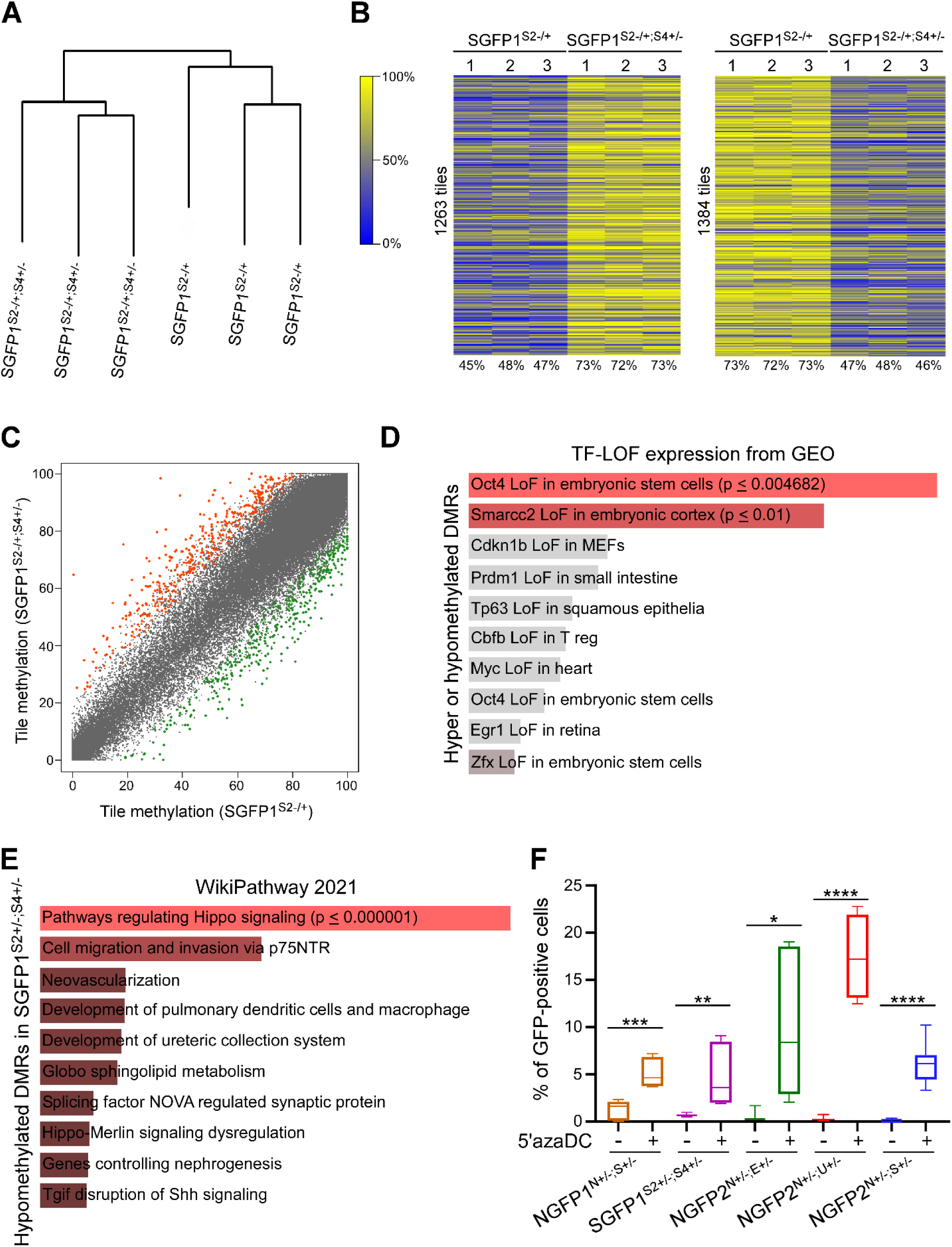
DNA methylation abnormalities in the double heterozygous mutant fibroblasts hinder the reprogramming process. **(A)** Dendrogram for SGFP1^S2+/-^ MEFs and SGFP1^S2+/-S4+/-^ MEFs based on the level of relative change observed at CpG sites with a threshold of 10 reads per site. **(B)** Heatmap of 20% of differentially methylated regions (DMRs) which covers a 100bp genomic region/tile and filtered to include at least 15 reads, which may be either hypermethylated or hypomethylated in SGFP1^S2+/-S4+/-^ MEFs compared to its parental SGFP1^S2+/-^ control MEFs. p-value ≤ 0.001. **(C)** Scatter plot analysis showing all the differentially methylated regions between SGFP1^S2+/-^ MEFs and SGFP1^S2+/-S4+/-^ MEFs (average of 3 biological replicates). Blue dots represent regions that are significantly more methylated in the double heterozygous mutant SGFP1^S2+/-S4+/-^ MEFs while red dots represent regions that are more methylated in the control SGFP1^S2+/-^ MEFs. Red dots represent regions that are associated with genes that are related to pluripotency and development and are significantly more methylated in SGFP1^S2+/-S4+/-^ MEFs while green dots represent regions that are associated with genes that are related to pluripotency and development and are significantly more methylated in SGFP1^S2+/-^ MEFs. Gray area represents no significant differences between the samples. **(D, E)** EnrichR of GEO (D) and wiki pathways analysis (E) of significantly over-represented genes that are either hyper or hypomethylated DMRs (D) or hypomethylated DMRs significantly enriched in SGFP1^S2+/-S4+/-^ MEFs (E) **(F)** Bar plot graph displaying the percentage of GFP-positive cells in the indicated samples after 13 days of reprogramming and 3 days of transgene removal with and without prior treatment of 5’azaDC for two days. Error bars indicate standard deviation between 6-7 biological replicates. *p-value ≤ 0.01, **p-value ≤0.001 as calculated by GraphPad Prism using 2-tailed Student’s t-test.

To confirm that DNA methylation abnormalities is responsible for the reprogramming delay seen in the double heterozygous mutant lines we next employed the DNA hypomethylation agent, the 5-Aza-2′-deoxycytidine (5’azaDC). Double heterozygous mutant fibroblasts from all lines were treated for two days with 5’azaDC and reprogramming experiments were carried out by the addition of dox for 13 days followed by 3 days of dox removal. In agreement with the RRBS results, treatment of 5’azaDC for two days prior to dox addition rescued the reprogramming defect seen in the double heterozygous mutant lines (**Figure 6F**). These results suggest that reduced levels of pluripotency genes at the pluripotent state leads to methylation abnormalities later on in their somatic cell derivatives. Moreover, these data suggest that the KI/KO approach to introduce a reporter gene should be avoided to maintain normal epigenetic state in the cells.

## DISCUSSION

Fluorescent reporter genes are a widely used tool in science to monitor the activity of a gene, regulatory element, non-coding RNA or other elements in the genome. One of the most common approach to introduce a reporter gene in a *locus-specific* manner is by the KI/KO approach. In this technique, the genomic sequence of the element of interest is being replaced by the coding sequence of the reporter gene, leaving only one intact allele of the targeted element. While this approach is considered harmless to the cells, up till now no thorough study has been conducted to support this claim. Here, by using pluripotent stem cells as a tested model we aimed to understand how allele elimination affects cell’s function and potential. To increase our ability to detect abnormalities caused by the elimination of the targeted alleles we deleted a single allele from various combinations of two pluripotency genes (i.e. Nanog^+/-^;Sall4^+/-^, Nanog^+/-^;Esrrb^+/-^, Nanog^+/-^;Utf1^+/-^, Sox2^+/-^;Sall4^+/-^) and used different pluripotent stem cell systems to exclude any system-specific effect. Interestingly, while examination of the developmental potential of the cells did not reveal a significant difference between the double heterozygous mutant cells and their parental controls, fibroblasts derived from these double heterozygous mutant pluripotent cells, demonstrated a strong blockage in their capability to induce pluripotency either by transcription factors or by nuclear transfer. The poor reprogramming efficiency observed between the various pluripotent stem cell systems ranged from a complete blockage at the mesenchymal to epithelial (MET) transition (NGFP2 line) to a later blockage at the stabilization step just before the acquisition of pluripotency (NGFP1 and SGFP1 lines, data not shown). Given that the affected genes were shown to play a major role during the stochastic phase of the reprogramming process (Buganim et al., 2012; Guo et al., 2019; Morshedi et al., 2013), we next examined the possibility that reduced stochastic expression of the targeted genes hinders the capability of the cells to pass the stochastic phase and to induce pluripotency. As the blockage seen in the double heterozygous mutant cells happened in the vast majority of the induced cells, to support the claim that reduced stochastic expression is responsible to the observed blockage, we aimed to show that the activation of the *Sall4* or *Nanog* allele is a frequent event and occurred in most induced cells at early stages of reprogramming. To test this hypothesis, we generated a tracing system for *Nanog* and *Sall4* that is based on the activity of Cre recombinase that unleashes an irreversible activation of a tdTomato reporter gene once activated. However, only a small number of induced cells turned on the tdTomato reporter following six days of factor induction, suggesting that reduced stochastic expression of these genes is not responsible for the global reprogramming delay seen in the double heterozygous mutant cells.

Additional expression of multiple pluripotent factors (e.g. *Sall4*, *Nanog*, *Utf1*, *Esrrb*, OSK) can either partially or fully rescue the observed blockage, thus, we next hypothesized that epigenetic barrier in the double heterozygous mutant fibroblasts, but not the allele elimination itself, may cause the observed delay. To that end, we subjected the parental and its double heterozygous mutant fibroblast counterparts to CpG-enriched DNA methylation analysis. A clear difference in the DNA methylation levels in regions within pluripotent and developmental genes was noted between the two fibroblast lines, suggesting that even a 50% reduction in the levels of two pluripotent genes is sufficient to induce aberrant DNA methylation during development. In fact, although *Oct4* expression level was not affected in the iPSCs, GEO enrichment of the derived MEFs showed loss of function of *Oct4* as a core pluripotency player which can be explained by the fact that key pluripotent genes such as *Nanog*, *Sox2*, *Sall4* and *Esrrb* who have been shown to regulate and function with the core DNA methylation machinery were missing (Adachi et al., 2018; Shanak and Helms, 2020; Tan et al., 2013).

The KI/KO approach is a well-accepted technique to introduce a reporter gene, however, there are other methodologies to introduce a reporter gene into a gene of interest without eliminating the gene’s coding sequences. This includes the use of self-cleavage peptides such as 2A and the internal ribosome entry site (IRES). Nevertheless, even in these, relatively safe techniques, in many occasions, a robust degradation of the targeted allele is still observed due to destabilization of the targeted mRNA following the introduction of the reporter cassette itself (Benchetrit et al., 2019). Overall, our study suggest that the KI/KO approach should be used carefully when cell state establishment is studied and emphasizes the importance of having two intact alleles for proper cellular functioning.

## Material and Methods

### Cell culture

Mouse embryonic fibroblasts (MEFs) were isolated as previously described (Wernig et al., 2008). MEFs were grown in DMEM supplemented with 10% fetal bovine serum, 1% non-essential amino acids, 2 mM L-Glutamine and antibiotics. ESCs and iPSCs were grown in S/Lif medium or 2i/Lif: DMEM supplemented with 10% fetal bovine serum, 1% non-essential amino acids, 2 mM L-Glutamine, 2X106 units mLif, 0.1 mM β-mercaptoethanol (Sigma) and antibiotics with or without 2i- PD0325901 (1 mM) and CHIR99021 (3 mM) (PeproTech). All the cells were maintained in a humidified incubator at 37°C and 6% CO2. All infections were performed on MEFs (passage 0-2) that were seeded at 50-70% confluency two days before the first infection. During the reprogramming to iPSC, the cells were grown in S/Lif medium with the addition of 2 μg/ml doxycycline.

### Secondary MEF production

Briefly, iPSC lines (NGFP2, NGFP1 and SGFP1 lines) were injected into blastocysts and chimeric embryos were isolated at E13.5.‏ For MEF production, embryos were dissected under the binocular removing internal organs and heads. The remaining body was chopped thoroughly by scalpels and exposed to 1ml Tripsin-EDTA (0.25%, GIBCO) for 30 minutes at 37°C. Following that, 10 mL of DMEM medium containing 10%FBS was added to the plate and the chopped tissue was subjected to thorough and intensive pipetting resulting in a relatively homogeneous mix of cells. Each chopped embryo was seeded in 15cm plate and cells were cultured in DMEM supplemented with 10%FBS, 2mM L-glutamine, and antibiotics until the plate was full. Puromycin (2 μg/ml) was added to each 15cm plate for positive selection for NGFP2, NGFP1 and SGFP1 MEFs, eliminating only the host cells.

### Immunostaining and Western blot

Cells were fixed in 4% paraformaldehyde (in PBS) for 20 minutes. The cells were rinsed 3 times with PBS and blocked for 1hr with PBS containing 0.1% Triton X-100 and 5% FBS. The cells were incubated overnight with primary antibodies (1:200) in 4C. The antibodies are: anti-SALL4 (Abcam, ab29112) and anti-NANOG (Bethyl, A300-379A) in PBS containing 0.1% Triton X-100 and 1%FBS (1:200 dilution). The next day, the cells were washed 3 times and incubated for 1hr with relevant (Alexa) secondary antibody in PBS containing 0.1% Triton X-100 and 1% FBS (1:500 dilution). DAPI (1:1000 dilution) was added 10 minutes before the end of incubation. For western blot, cell pellets were lysed on ice in lysis buffer (20 mM Tris-HCl, pH 8, 1mM EDTA pH 8, 0.5% Nonidet P-40, 150mM NaCl, 10% glycerol, 1mM, protease inhibitors (Roche Diagnostics) for 10 min, supernatant were collected and 40μg protein were suspended with sample buffer and boiled for or 5 min at 100C, and subjected to western blot analysis. Primary antibodies: anti-SALL4 (Abcam, ab29112), anti-NANOG (Bethyl, A300-379A), anti-ESRRB (Perseus proteomics, PP-H6705-00), anti-UTF1 (Abcam, ab24273), anti-Actin (Santa cruz, sc-1616), anti-β-Tubulin (Abcam, ab179513), anti-Vinculin (Abcam, ab129002). Blots were probed with anti-mouse, anti-goat or anti-rabbit IgG-HRP secondary antibody (1:10,000) and visualized using ECL detection kit.

### Southern Blot

Southern blot was performed as previously described (Carey et al., 2011).

### FACS analysis

Cells were washed twice with PBS and trypsinized (0.25%) and filtered through mesh paper. Flow cytometry analysis was performed on a Beckman Coulter and analyzed using Kaluza Software. All FACS experiments were repeated at least three times, and the bar graph results are presented as a mean ± standard deviation of two biological duplicate from a typical experiment. Flow cytometry analysis was performed on a Beckman Coulter and analyzed using Kaluza Software.

### Quantitative real-time PCR

Total RNA was isolated using the Macherey-Nagel kit (Ornat). 500–2000 ng of total RNA was reverse transcribed using iScript cDNA Synthesis kit (Bio-Rad). Quantitative PCR analysis was performed in duplicates using 1/100 of the reverse transcription reaction in a StepOnePlus (Applied Biosystems) with SYBR green Fast qPCR Mix (Applied Biosystems). Specific primers flanking an intron were designed for the different genes (see Primer Table). All quantitative real-time PCR experiments were repeated at least three times, and the results were normalized to the expression of *Gapdh* and presented as a mean ± standard deviation of two duplicate runs from a typical experiment.

### RNA sequencing

Total RNA was isolated using Rneasy Kit (QIAGEN) and sent to the “Technion Genome Center”, Israel, for library preparation and sequencing.

### Cleaning and filtering of raw reads

Raw reads (fastq files) were inspected for quality issues with FastQC (v0.11.2, http://www.bioinformatics.babraham.ac.uk/projects/fastqc/). According to the FastQC report, reads were then trimmed to a length of 50 bases with fastx_trimmer of the FASTX package (version 0.0.13, http://hannonlab.cshl.edu/fastx_toolkit/), and quality-trimmed at both ends, using in-house perl scripts, with a quality threshold of 32. In short, the scripts use a sliding window of 5 base pairs from the read’s end and trim one base at a time until the average quality of the window passes the given threshold. Following quality-trimming, adapter sequences were removed by Trim Galore (version 0.3.7, http://www.bioinformatics.babraham.ac.uk/projects/trim_galore/), using the command “trim_galore -a $adseq –length 15” where $adseq is the appropriate adapter sequence. The remaining reads were further filtered to remove very low-quality reads, using the fastq_quality_filter program of the FASTX package, with a quality threshold of 20 at 90 percent or more of the read’s positions.

### Expression analysis

The cleaned fastq files were mapped to the mouse transcriptome and genome, Ensembl version GRCm38 from Illumina’s iGenomes (http://support.illumina.com/sequencing/sequencing_software/igenome.html), using TopHat (v2.0.11), allowing up to 3 mismatches and a total edit distance of 8 (full command: tophat -G Mus_musculus/Ensembl/GRCm38/Annotation/Genes/genes.gtf -N 3 --read-gap-length 5 --read-edit-dist 8 --segment-length 18 --read-realign-edit-dist 5 --b2-i S,1,0.75 --b2-mp 3,1 --b2-score-min L,-0.5,-0.5 Mus_musculus/Ensembl/GRCm38/Sequence/Bowtie2Index/genome clean.fastq). Quantification and normalization were done with the Cufflinks package (v2.2.1). Quantification was done with cuffquant, using the genome bias correction (-b parameter), multi-mapped reads assignment algorithm (-u parameter) and masking for genes of type IG, TR, pseudo, rRNA, tRNA, miRNA, snRNA and snoRNA (-M parameter). Normalization was done with cuffnorm (using output format of Cuffdiff).

### Visualization

The R package cummeRbund (version 2.8.2) was used to calculate and draw the figures (such as scatter plots, MA plots, etc.) from the normalized expression values.

### Chimera Formation

Blastocyst injections were performed using (C57/Bl6xDBA) B6D2F2 or CB6F1 host embryos. All injected iPSC lines were derived from crosses of 129Sv/Jae to C57/Bl6 mice and could be identified by agouti coat color. Embryos were obtained 24 hr (1 cell stage) or 40 hr (2 cell stage) posthuman chorionic gonadotropin (hCG) hormone priming. Diploid embryos were cultured in EmbryoMax KSOM (Millipore) or Evolve KSOMaa (Zenith Biotech) until they formed blastocysts (94–98 hr after hCG injection) at which point they were placed in a drop of Evolve w/HEPES KSOMaa (Zenith) medium under mineral oil. A flat tip microinjection pipette with an internal diameter of 16 mm (Origio) was used for iPSC injections. Each blastocyst received 8–12 iPSCs. Shortly after injection, blastocysts were transferred to day 2.5 recipient CD1 females (20 blastocysts per female). Pups, when not born naturally, were recovered at day 19.5 by cesarean section and fostered to lactating Balb/c mothers.

### Nuclear transfer

Nuclear transfer was performed as described (Wakayama et al., 1998) with modifications. Briefly, metaphase II-arrested oocytes were collected from superovulated B6D2F1 females (8-10 wks) and cumulus cells were removed using hyaluronidase. The oocytes were enucleated in a droplet of HEPES-CZB medium containing 5μg/ml cytochalasin B (CB) using a blunt Piezo-driven pipette. After enucleation, the spindle-free oocytes were washed extensively and maintained in CZB medium up to 2 h before nucleus injection. The CCs from mice (B6D2F1) were aspirated in and out of the injection pipette to remove the cytoplasmic material and then injected into enucleated oocytes. The reconstructed oocytes were cultured in CZB medium for 1 h and then activated for 5-6 h in activation medium containing 10mM Sr 2+, 5ng/ml trichostatin A (TSA) and 5μg/ml CB. Following activation, all of the re constructed embryos were cultured in KSOM medium supplemented with 5ng/ml TSA for another 3-4 hours and maintained in KSOM medium with amino acids at 37°C under 5% CO2 in air.

### Reduced-representation bisulfite sequencing (RRBS)

DNA was isolated from MEFs and incubated in lysis buffer (25 mM Tris-HCl (pH 8), 2 mM ethylenediaminetetraacetic acid, 0.2% sodium dodecyl sulfate, 200 mM NaCl) supplemented with 300 μg/mL proteinase K (Roche) followed by phenol:chloroform extraction and ethanol precipitation and RRBS libraries were prepared (Boyle et al., 2012) and run on HiSeq 2500 (Illumina) using 100 bp paired-end sequencing.

DNA methylation was analyzed by using 100 bp paired-end sequencing reads from RRBS that were trimmed and quality filtered by trim galore software using default parameters for RRBS. Read alignment (genome build mm10) and extraction of single-base resolution methylation levels were carried out by BSMAP. Differentially methylated regions (DMR) were explored with R methylKit package version 1.18.0 (Akalin et al., 2012). CpG sites featuring less than 10 reads were considered unreliable and discarded from further analysis. CpG sites were then aggregated into consecutive tiles of size 100 bp and a threshold of at least 15 reads per tile was applied. Differential methylation between the two lines, each consisting of three samples, was determined by logistic regression and adjusted p-values are calculated with SLIM (sliding linear model). A threshold of 1E-3 was set for adjusted p-value and a threshold of 20 methylation points was set between the two lines and further explored. DMRs were annotated with Homer (Hypergeometric Optimization of Motif Enrichment) version 4.11.1 (Heinz et al., 2010) and specifically its function annotatePeaks.pl. This function outputs a set of genes affiliated with DMR based on the nearest promoter distance. Heatmaps were created with R package heatmap.2 version 3.1.1 and dendrogram with R package dendextend version 1.15.2 (Galili, 2015).

## ACKNOWLEDGEMENTS

Y.B. is supported by research grants from EMBO Young Investigator Programme (YIP), DKFZ-MOST (CA 177), Howard Hughes Medical Institute International Research Scholar (HHMI, #55008727) and by a generous gift from Ms. Nadia Guth Biasini. We thank Yuval Nevo and huji core bioinformatics unit for analyzing part of the RNA-seq data.

## Author Information

RNA-seq data for the various NGFP2^N+/-^ double heterozygous mutant and control iPSC lines (accession number GSE182009) and RNA-seq for NGFP2^N+/-^ double heterozygous mutant and control MEF lines after 6 days of reprogramming and RRBS for the SGFP1^S2+/-^ and SGFP1^S2+/-;S4+/-^ primary MEFs (accession number GSE192655) has been deposited to the Gene Expression Omnibus database (GEO). The reviewers can enter the deposited data using the following token: wxcbcokaplurhsn. Correspondence and requests for materials should be addressed to Y.B.

## Author Contribution

Y.B. and R.J. conceived the study. Y.B., R.L. designed the experiments, prepared the figures and wrote the manuscript. Y.B. together with E.K., C.O., and D.F. generated the NGFP2^N+/-^ double heterozygous mutant lines and ran the various reprogramming experiments on NGFP2^N+/-^ lines. R.L. generated the tracing systems, with the help of N.M. for Nanog and Sall4 and the NGFP1^N+/-;S+/-^, NGFP1^N-/-^ and SGFP1^N+/-;S4+/-^ lines, performed reprogramming experiments on these lines, immunostaining, flow cytometry and 5’azaDC experiments. N.M. prepared the samples for the RNA-seq at day 6 of reprogramming and performed qPCR for the MET genes. N.M. together with N.YT. ran rescue reprogramming experiments and performed sm-mRNA-FISH. A.W.C. analyzed the RNA-seq data from the various NGFP2^N+/-^ iPSC lines. H.Y. performed SCNT experiments. S.M. and K.M. injected iPSC lines to produce secondary MEFs and chimeric mice. M.A. helped R.L. to run reprogramming experiments and to analyze the flow cytometry results.

---

**Figure S1.**
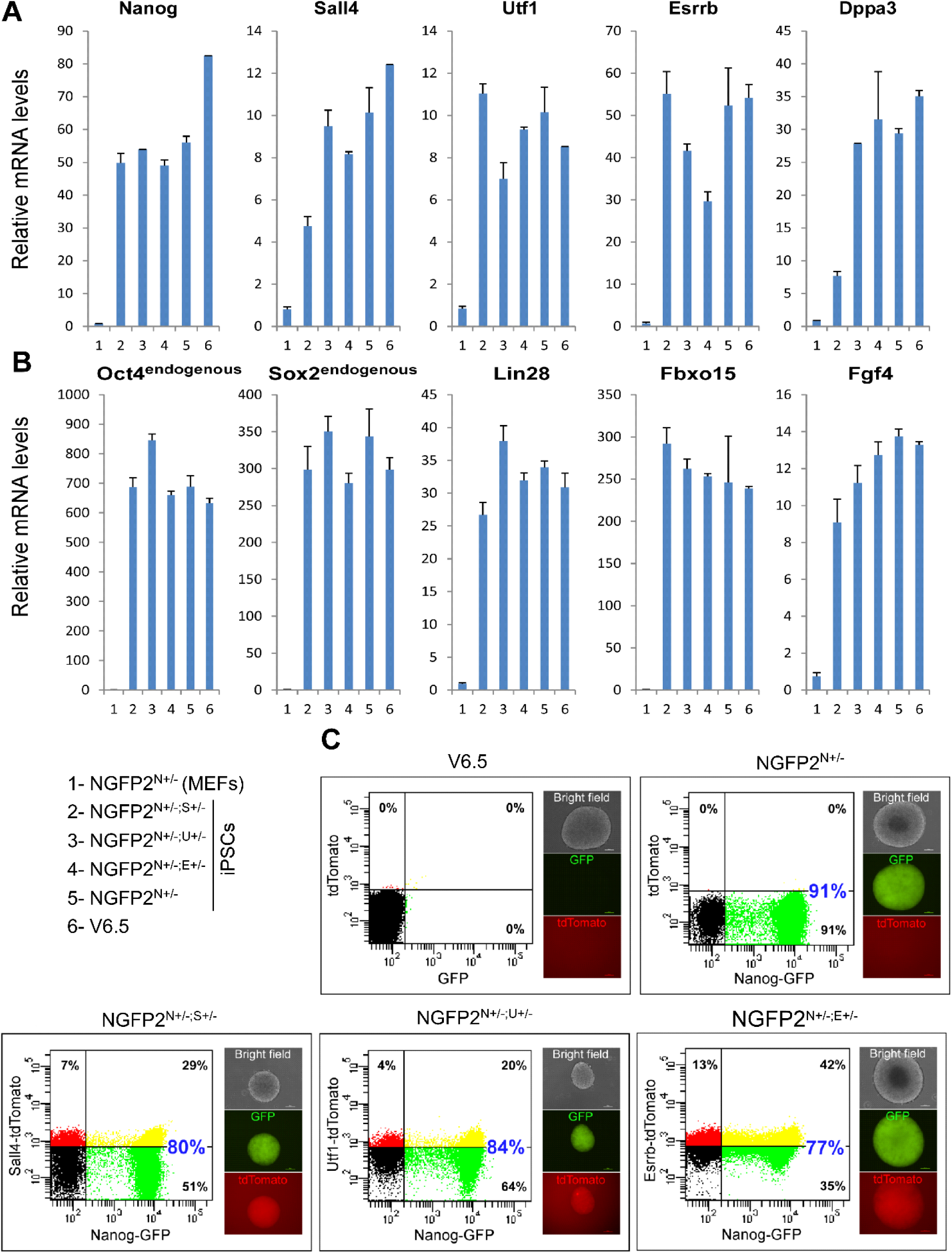
Characterization of the double heterozygous mutant NGFP2^N+/-^ lines. **(A)** qPCR of the indicated genes normalized to the housekeeping control gene *Gapdh* in the various NGFP2^N+/-^ double heterozygous mutant lines, NGFP2^N+/-^ parental line, and ESC (V6.5) and MEF controls. Error bars presented as a mean ± SD of 2 duplicate runs from a typical experiment out of 3 independent experiments. **(B)** Flow cytometry analysis for GFP (*Nanog*) and tdTomato (*Utf1, Esrrb* or *Sall4*) in the various double heterozygous mutant lines that grew under 2i/Lif conditions. Note that although tdTomato lacks polyA, a red signal is still detectable

**Figure S2.**
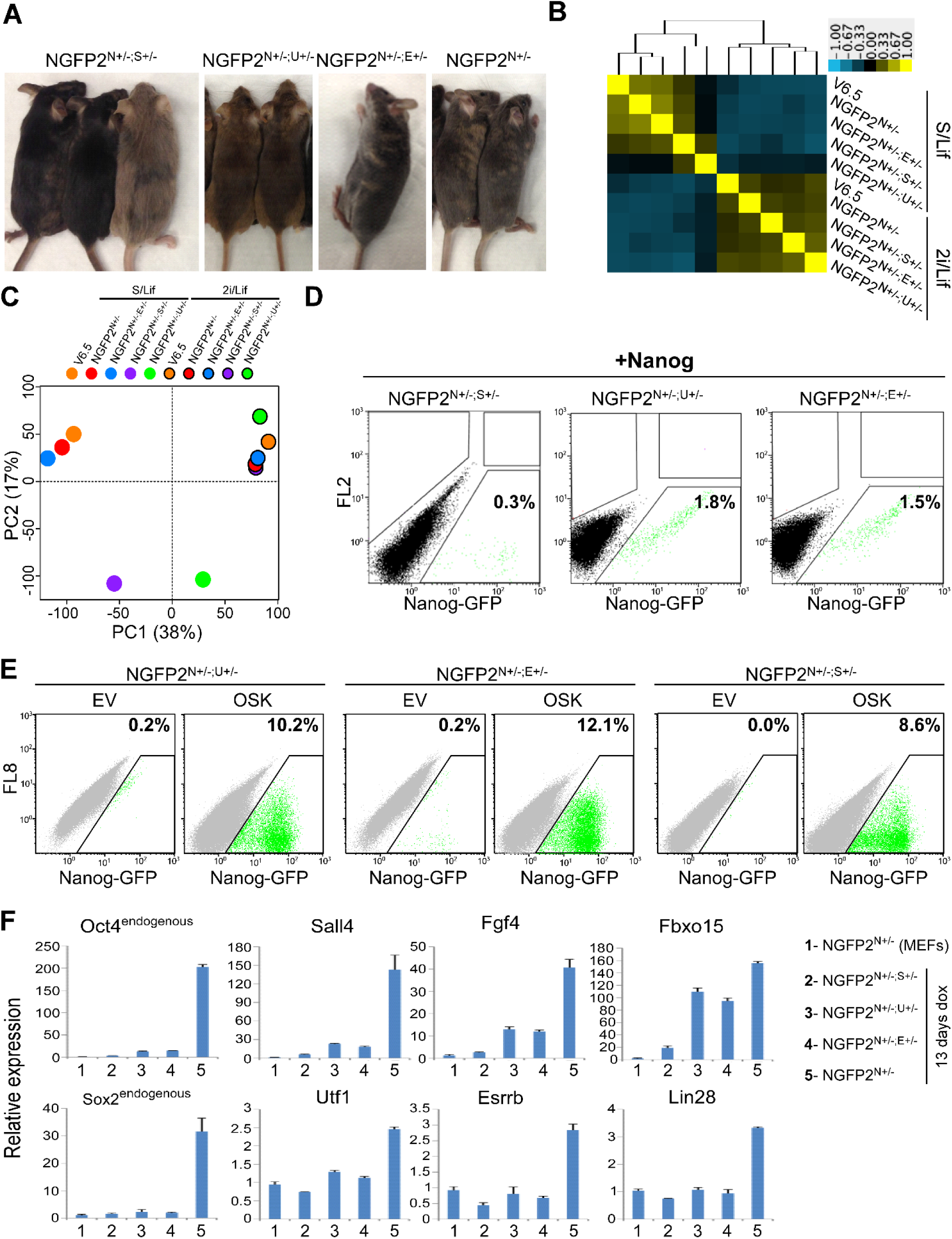
The developmental potential and transcriptional profile of NGFP2^N+/-^ double heterozygous mutant lines and rescue reprogramming experiments. **(A)** Representative images of adult chimeric mice produced by the various NGFP2^N+/-^ double heterozygous mutant iPSC lines and control following blastocyst injection and transplantation into foster mothers. **(B)** Correlation heatmap and dendrogram of global gene expression profiles for two RNA-seq replicates for the indicated NGFP2^N+/-^ iPSC lines and ESC (V6.5) control grown under S/Lif or 2i/Lif conditions. Replicate pairs were assigned a shared numerical value. **(C)** Principle component analysis for the indicated samples using 500 most differentially expressed genes among all samples. PC1, 38%; PC2, 17%. Each line is marked by a specific color. The group names correspond to the names in (B). Cells that were grown in 2i/Lif are surrounded with black circle. **(D, E)** Flow cytometry analysis of Nanog-GFP-positive cells for the various NGFP2^N+/-^ double heterozygous mutant lines following overexpression of *Nanog* (D) or OSK (E) and reprogramming for 13 days following by 3 days of dox removal. OSK indicates *Oct4*, *Sox2* and *Klf4* and EV indicates empty vector **(F)** NGFP2^N+/-^ double heterozygous mutant lines and control were reprogrammed for 13 days. qPCR analysis showing the expression levels of the indicated genes, in the depicted samples, after *Gapdh* normalization. Error bars presented as a mean ± SD of 2 duplicate runs from a typical experiment out of 3 independent experiments.

**Figure S3.**
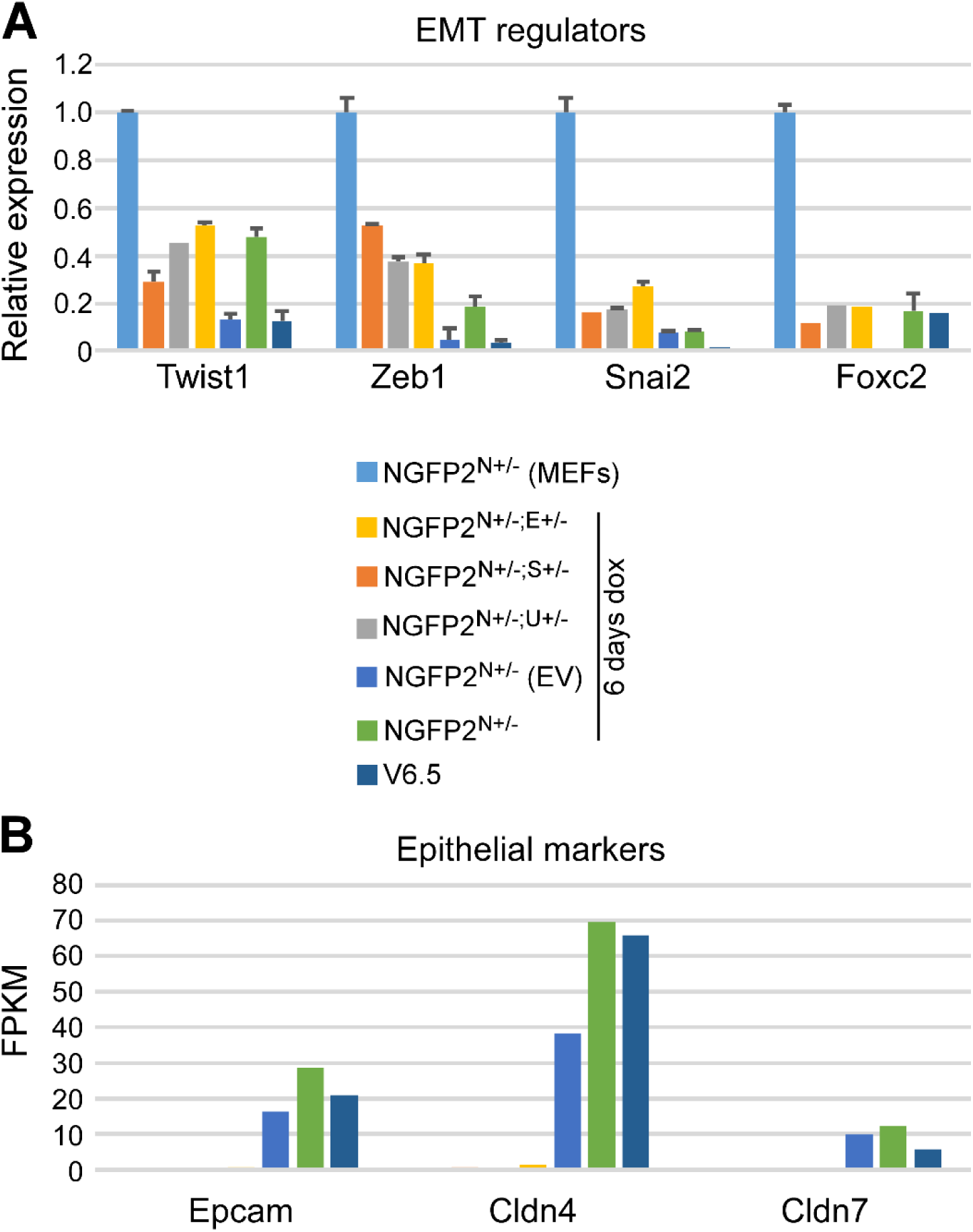
NGFP2^N+/-^ double heterozygous mutant lines fail to activate the epithelial program during reprogramming. **(A)** qPCR of the indicated EMT genes normalized to housekeeping control gene *Gapdh* in the various NGFP2^N+/-^ double heterozygous mutant lines following 6 days of dox and in ESCs (V6.5) and NGFP2^N+/-^ MEF control. Error bars presented as a mean ± SD of 2 duplicate runs from a typical experiment out of 3 independent experiments. **(B)** Graph summarizing the expression level (FPKM-Fragments Per Kilobase Million) of the indicated epithelial genes in the various NGFP2^N+/-^ double heterozygous mutant lines after 6 days of dox and in ESCs (V6.5) and NGFP2^N+/-^ MEF control. Expression level of the depicted genes was obtained from the RNA-seq data described in Figure 3.

**Figure S4.**
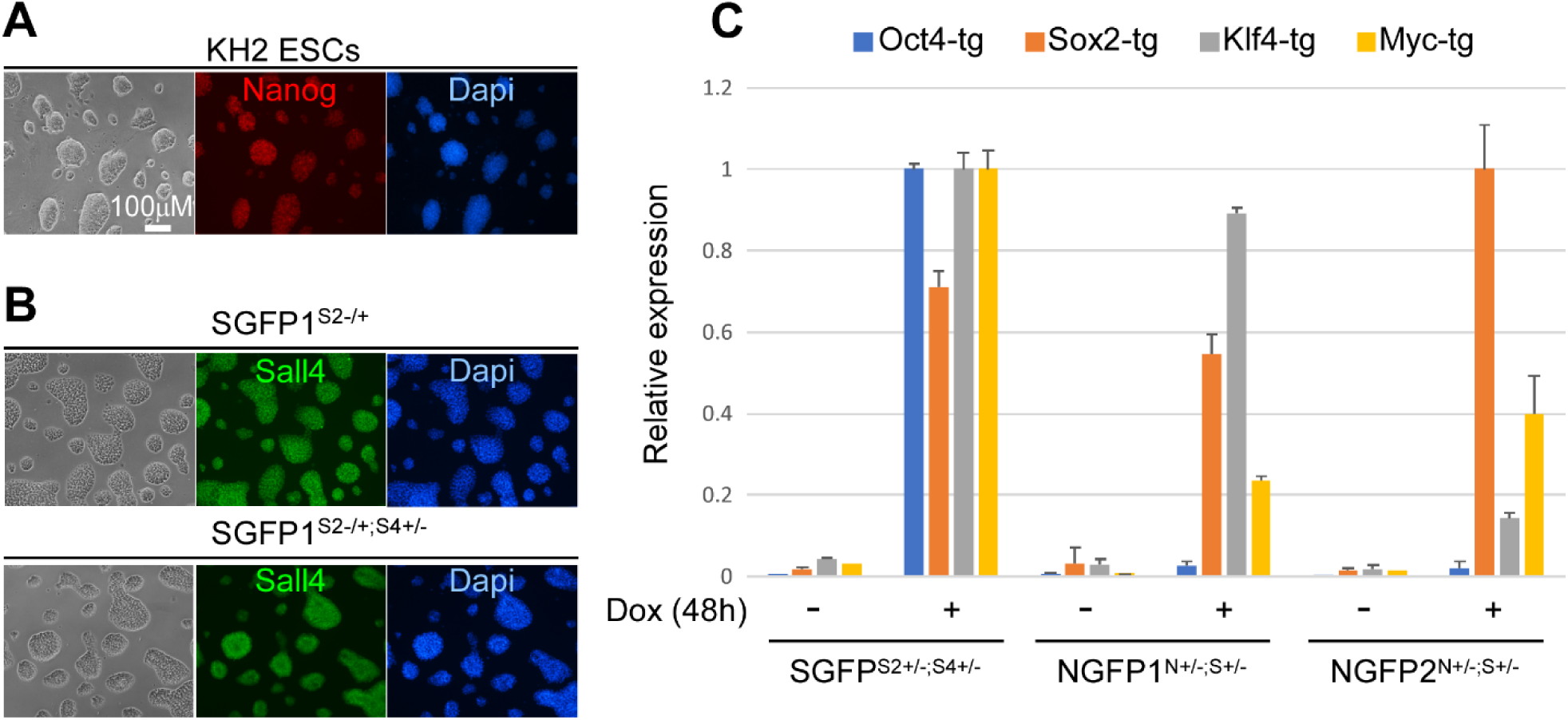
Nanog and Sall4 protein level in targeted iPSC lines and controls. **(A)** Bright field images and immunostaining images for *Nanog* (red) and Dapi (blue) in KH2 ESCs. Scale bar, 100μM. **(B)** Bright field images and immunostaining images for Sall4 (green) and Dapi (blue) in SGFP1^S2+/-^ and SGFP1^S2+/-; S4+/-^ iPSC lines. Scale bar, 100 uM. **(C)** qPCR of the indicated OSKM transgenes normalized to housekeeping control gene *Gapdh* in the various double heterozygous mutant MEF lines following 2 days of culture with or without dox. Error bars presented as a mean ± SD of 2 duplicate runs from a typical experiment out of 3 independent experiments.

**Figure S5.**
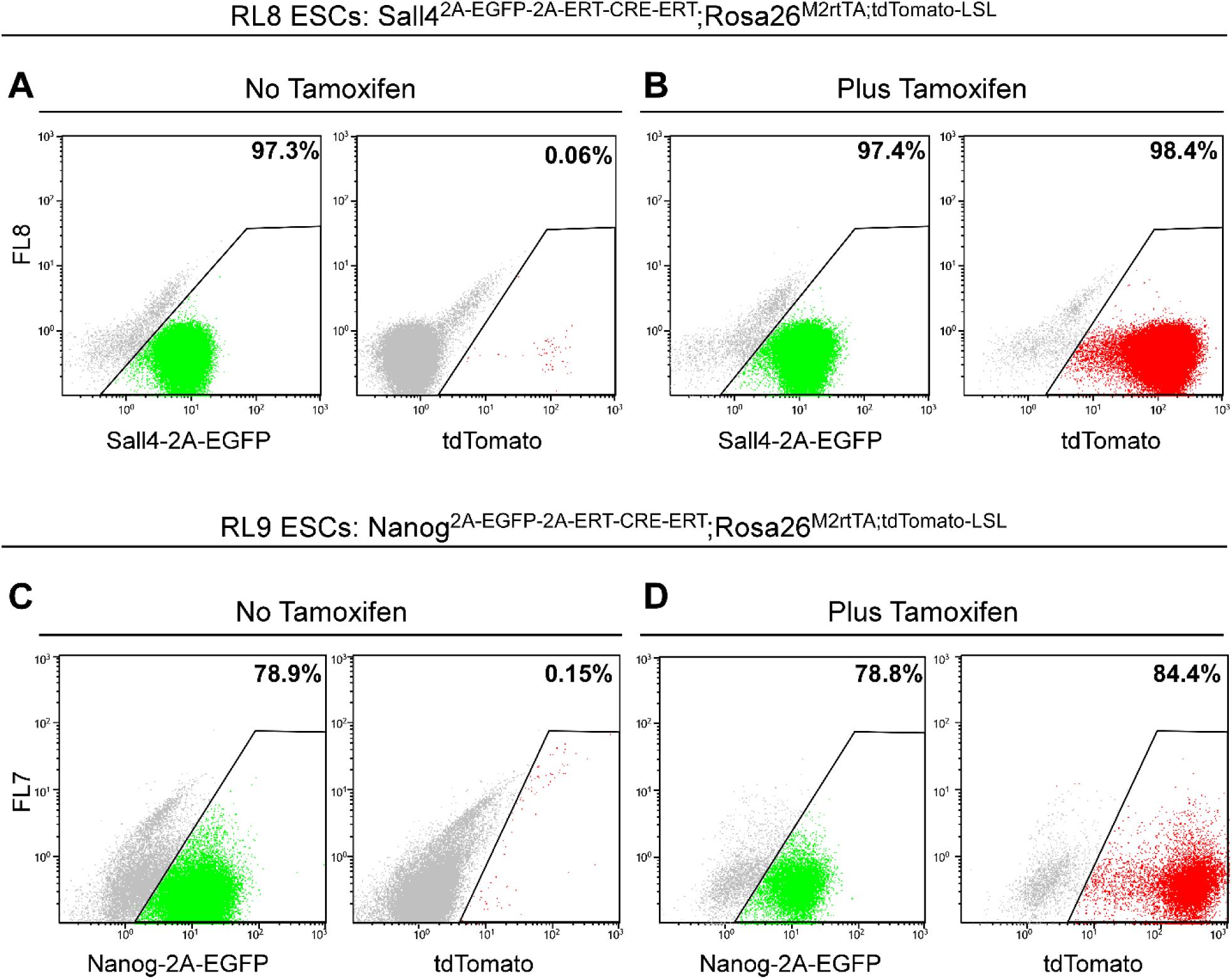
Sall4 and Nanog tracing system characterization. **(A)** Flow cytometry analysis for Sall4-2A-EGFP and tdTomato in the targeted ESC clone RL8 before and after tamoxifen addition (48 hours). **(B)** Flow cytometry analysis for Nanog-2A-EGFP and tdTomato in the targeted ESC clone RL9 before and after tamoxifen addition (48 hours).

**Table 1.**
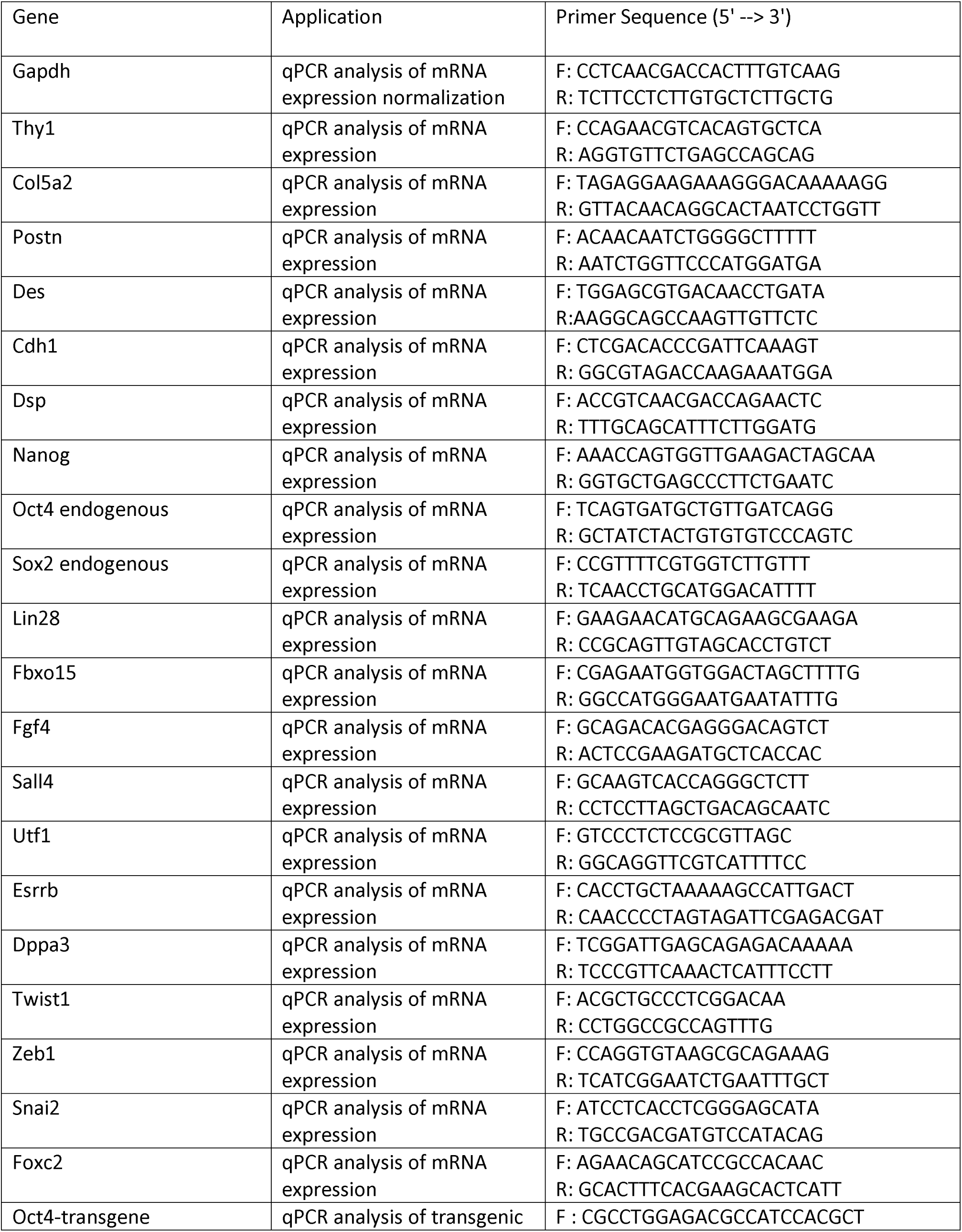
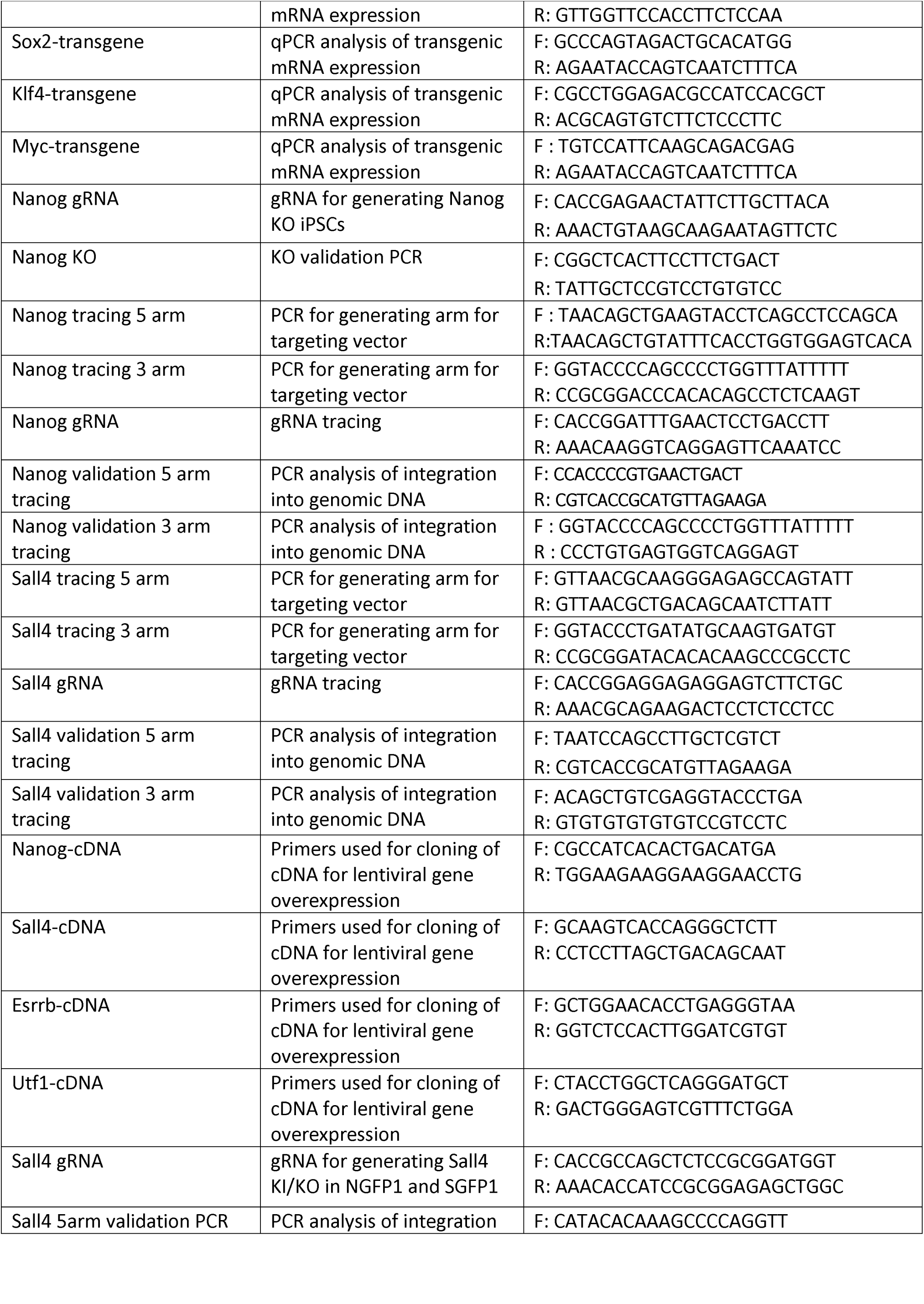
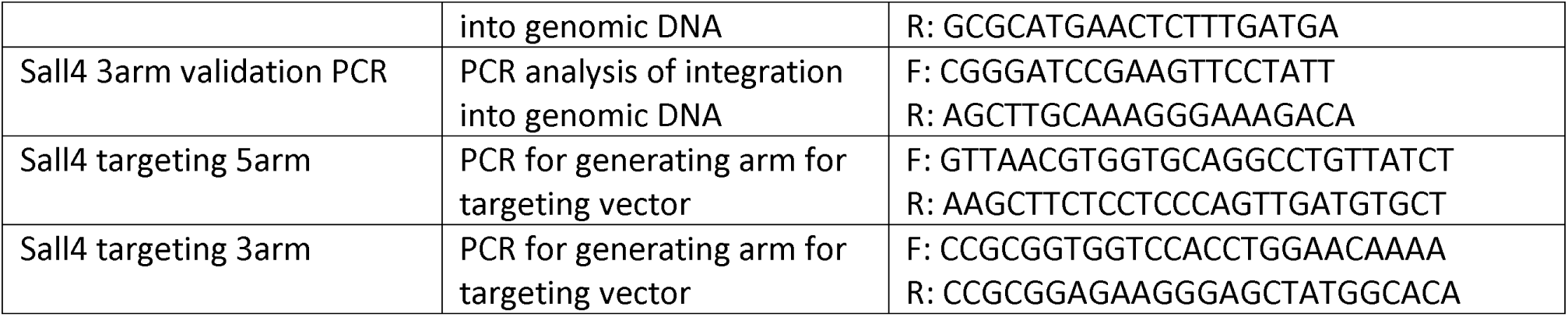
primer list

